# Structural Determinants for Activity of the Antidepressant Vortioxetine at Human and Rodent 5-HT_3_ receptors

**DOI:** 10.1101/2024.02.23.581731

**Authors:** Uriel López-Sánchez, Lachlan Jake Munro, Lucy Kate Ladefoged, Anders Juel Pedersen, Christian C. Nielsen, Signe M. Lyngby, Delphine Baud, Sarah C. R. Lummis, Benny Bang-Andersen, Birgit Schiøtt, Christophe Chipot, Guy Schoehn, Jacques Neyton, Francois Dehez, Hugues Nury, Anders S. Kristensen

## Abstract

Vortioxetine (VTX) is a recent antidepressant that targets a variety of serotonin receptors. We investigate the drug’s molecular mechanism of operation at serotonin 5-HT_3_ receptors (5-HT_3_R), which features two mysterious properties: VTX acts differently on rodent and human 5-HT_3_R; VTX appears to suppress any subsequent response to agonists. Using a combination of cryo-EM, electrophysiology, and molecular dynamics, we show that VTX stabilizes a resting inhibited state of the mouse 5-HT_3_R and an agonist bound-like state of the human 5-HT_3_R, in line with the functional profile of the drug. We report four human 5-HT_3_R structures and show that the human receptor transmembrane domain is intrinsically fragile. We also explain the lack of recovery after VTX administration via a membrane partition mechanism.

## Introduction

Vortioxetine (VTX) is a recently approved antidepressant drug for treating major depressive disorder. Unlike the selective-serotonin reuptake inhibitor (SSRI) class of antidepressants that selectively target the serotonin transporter (SERT), VTX has a multimodal activity profile across the serotonergic system with affinity for the SERT and a range of serotonin receptors that include the metabotropic 5-HT_1A_, 5-HT_1B_, 5-HT_1D_ and 5-HT_7_ receptors and the ionotropic 5-HT_3_ receptors (Bang-Andersen et al., 2011; Sanchez et al., 2015). The metabotropic 5-HT receptor activity of VTX has been shown to contribute, in animal models, to the antidepressant effects independent of SERT (Nackenoff et al., 2017). The activity at the 5-HT_3_ receptor may contribute to advantageous properties of VTX observed in some pre-clinical models and clinical trials compared to SSRI and other monoamine transporter selective antidepressants (Mahableshwarkar et al., 2016; McIntyre et al., 2014; Pehrson et al., 2015; Riga et al., 2016).

In contrast to SERT and metabotropic 5-HT receptors, the mechanism of VTX at the ionotropic 5-HT_3_ receptors is not well understood. 5-HT_3_ receptors are pentameric ligand-gated ion channels (pLGICs) (Barnes et al., 2021; Nemecz et al., 2016). Like every member of the pLGIC family, functional 5-HT_3_ receptors contain five subunits. The homopentameric 5-HT_3A_ receptor (5-HT_3A_R) is the most studied subtype, yet heteromeric subtypes exist (Niesler, 2011). A 5-HT_3A_R subunit comprises three domains: an extracellular domain (ECD), a transmembrane domain (TMD) harboring the cationic pore, and an intracellular domain (ICD). VTX binds with low nanomolar affinity to the orthosteric binding sites located in clefts at the ECD subunit-subunit interfaces (Ladefoged et al., 2018). For pLGICs in general, extensive work has focused on understanding the conformational transitions that underlie activation and desensitization (Colquhoun and Lape, 2012; Gielen and Corringer, 2018). In recent years, an avalanche of pLGIC structures has provided a framework to interpret the wealth of functional results accumulated over decades of research in a structural context (Kim et al., 2020; Kumar et al., 2020; Masiulis et al., 2019; Yu et al., 2019). Specifically, structures of pLGICs captured in different functional states helped define common key mechanistic features, such as the dewetting of the upper pore in the resting and inhibited states. For 5-HT_3_ receptors, structural studies initiated by a crystal structure of a closed state (Hassaïne et al., 2014) now cover a range of conformations assigned to inhibited/resting, pre-active and open states (Basak et al., 2018; Polovinkin et al., 2018; Zhang et al., 2021). Structures are also available for complexes with antiemetic drugs of the -setron class, the major class of clinical drugs targeting the 5-HT_3_ receptor, which act as competitive antagonists to alleviate the adverse effects of chemotherapies (Basak et al., 2019; Zarkadas et al., 2020). Together, those structures revealed the precise geometry of the ECD and the TMD and how agonist versus antagonist-bound structures differ through reorganizing the ECD quaternary structure. However, no structure has yet been unambiguously assigned to the desensitized state, and the location of the gate obstructing the ion flux in the desensitized state remains elusive. In addition, only mouse 5-HT_3A_ receptor structures have been obtained to date.

At the human homomeric 5-HT_3A_R (h5-HT_3A_R), VTX is at present best described as an atypical ligand as it appears to have both agonistic and antagonistic effects. Specifically, VTX binding induces initial activation followed by a persistent and insurmountable inhibition of subsequent stimulation by serotonin (Bang-Andersen et al., 2011; Dale et al., 2018; Ladefoged et al., 2018). Thus, VTX acts as a functional antagonist under steady-state conditions. This mode of action has been hypothesized to reflect the ability of VTX to induce and stabilize a non-conducting desensitized state of the h5-HT_3A_R (Bang-Andersen et al., 2011; Dale et al., 2018). Desensitization is a hallmark functional feature of virtually every pLGIC and follows the transient opening of the channel upon prolonged application of an agonist (Gielen and Corringer, 2018). Previous work has suggested that the rate of VTX-induced desensitization in human receptors is an order of magnitude faster than serotonin and other synthetic agonists. In addition, the recovery from VTX-induced desensitization appears extremely slow (Bang-Andersen et al., 2011; Dale et al., 2018; Ladefoged et al., 2018). Another enigmatic aspect of VTX is its effects markedly differ across 5-HT_3A_ receptors from different mammalian species. Specifically, the agonist activity observed at the h5-HT_3A_R is absent for the rat 5-HT_3A_ receptor (r5-HT_3A_R) (Bang-Andersen et al., 2011). Additionally, the steady-state inhibitory potency (IC_50_) of VTX varies by more than three orders of magnitude between r5-HT_3A_R, h5-HT_3A_R, and the equivalent guinea pig receptor (gp5-HT_3_R) (Bang-Andersen et al., 2011; Dale et al., 2018).

In the present study, we seek to characterize VTX activity at human and rodent 5-HT_3A_Rs in further detail and, most importantly, identify and understand the unknown molecular determinants for the intriguing properties of VTX. First, we confirm and elaborate on the differential activity of VTX at human and rodent 5-HT_3A_ receptors. Second, we use cryo-electron microscopy (cryo-EM) and molecular dynamics (MD) to show that VTX stabilizes a resting-like inhibited state in the m5-HT_3A_R, similar to those of antiemetic-bound receptors. In contrast, the h5-HT_3A_R structure in complex with VTX features an ECD conformation similar to agonist-bound structures, in line with VTX stabilizing the desensitized state. We substantiate this structural observation by studying the effect of VTX on TMD mutants with accelerated desensitization and by monitoring local changes at the pore level with voltage-clamp fluorometry (VCF). We identify the protein residues that determine species-specific properties and take advantage of a series of VTX analogs to refine the drug structure-activity relationship. Finally, we provide evidence suggesting membrane lipophilic partitioning explains the persistent inhibition following long VTX applications.

## Results

### VTX is an agonist at the human 5-HT3AR but not at rodent receptors

Building on the previous work that suggests VTX acts as an agonist at human 5-HT_3A_ receptors but as an antagonist at rodent receptors (Bang-Andersen et al., 2011; Dale et al., 2018), we first characterized the agonist activity of VTX at 5-HT_3A_Rs from human, rat, mouse, and guinea pig (Fig. 1). For the human receptor expressed in *Xenopus* oocytes, application of 10 µM VTX or 300 µM serotonin evoked inward currents that reached peak amplitude within 1-2 s and then strongly decayed, consistent with a desensitization process (Fig. 1B). The peak-current amplitude of 10 µM VTX was 55±5 % (n=31) of the currents evoked by 300 µM serotonin (Table SI1). For both ligands, the current decays followed a monoexponential time course with a rate of 2.3±0.8 s (n=5) for VTX compared to 76±18 s (n=5) for serotonin (Fig. 1D). The EC_50_ for VTX activation was determined from dose-response curves to be 0.15 μM (0.10;0.25, n=4) for VTX compared to 1.90 μM (0.10;0.25, n=4) for serotonin (Fig. 1C). The serotonin-evoked currents from r5-HT_3A_R and mouse receptors (m5-HT_3A_R) displayed desensitization akin to that of h5-HT_3A_R, with no or barely detectable current remaining at steady-state conditions. Guinea pig receptors (gp5-HT_3A_R) did not desensitize fully, resulting in approximately 50% of the peak current remaining at steady-state conditions (Fig. 1B). For all three rodent receptors, VTX in concentrations up to 100 µM yielded no responses, confirming that VTX does not act as an agonist at these species (Fig. 1B). The potency of VTX at steady-state inhibition has previously been reported to vary dramatically between the guinea pig receptor and the human, rat and mouse receptors with the IC_50_ at gp5-HT_3A_R (720 nM) being two orders of magnitude higher than at r5-HT_3A_R (IC_50_ = 0.18 nM), m5-HT_3A_R (IC_50_ = 3.3 nM), h5-HT_3A_R (IC_50_ = 10 nM) (Dale et al., 2018). Using a membrane-potential fluorescent assay, we determined concentration-inhibition relationships of VTX for inhibition of serotonin activity and observed a similar rank-order of VTX IC_50_ as r5-HT_3A_R<h5-HT_3A_R<< gp5-HT_3A_R (Table SI1, SI2, Fig. 1e). Notably, the classical setron antagonists granisetron and ondansetron displayed similar rank-order of inhibitory potency (Table SI2). To characterize the recovery from VTX inhibition across the different receptors, we next employed a 3-step electrophysiological recording protocol, with an initial application of 10 µM 5-HT, followed by a super-saturating concentration of 10 µM VTX, and a final second application of 10 µM 5-HT with 6-minute wash periods in between (Fig. 1F). Following wash-out, h5-HT_3_R showed no detectable subsequent serotonin current. The r5-HT_3_R did show a small but detectable current of around 2%, whereas m5-HT_3A_R displayed a partial current recovery of around 50%. In contrast, gp5-HT_3A_Rs fully recovered from VTX inhibition (Table SI1). For h5-HT_3_R, we further tracked the lack of recovery and found full inhibition to persist for up to 25 minutes (Fig. 1G). Altogether, VTX acts as an antagonist at rodent receptors but as a fast-desensitizing agonist at human receptors in TEVC protocols. In addition, VTX displays significant species differences in inhibitory potency and rate of recovery from inhibition.

**Fig. 1:**
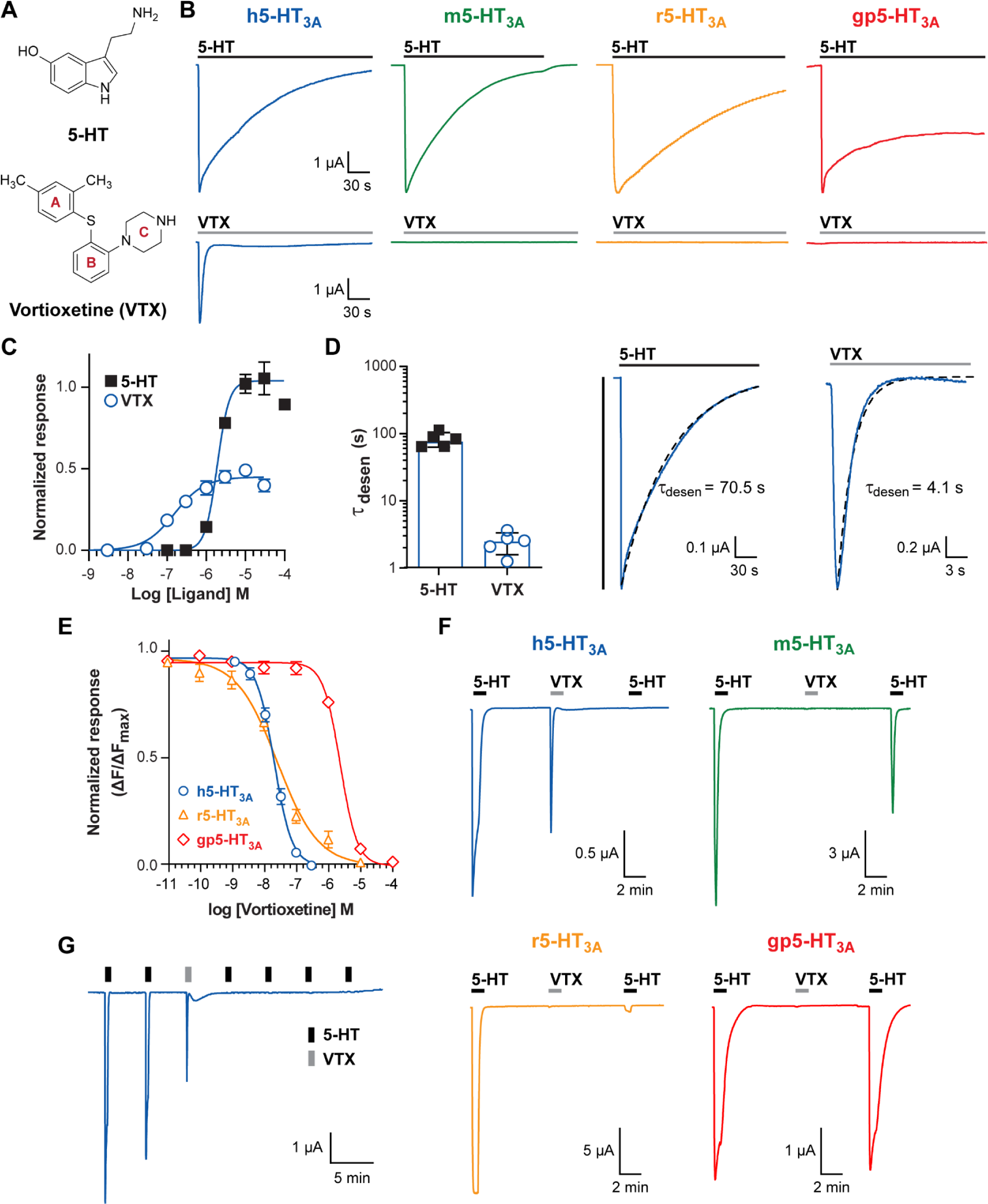
VTX acts as a fast-desensitizing, partial agonist at human receptors, and as an antagonist at rodent receptors. **A.** Chemical structure of the endogenous ligand serotonin (5-HT) and of the antidepressant vortioxetine (VTX). **B.** Representative traces illustrating receptor current activation and desensitization during application of 5-HT (300 µM) and VTX (10 µM) to *Xenopus* oocytes expressing h5-HT3A, m5-HT3A r5-HT3A and gp5-HT3A as indicated. **C.** 5-HT (filled squares) and VTX (circles) concentration response curves in *Xenopus* oocytes expressing h5-HT3A, receptors. Data points represent the mean ± SEM from at least 5 individual oocytes. **D.** Rate of desensitization (τdesen) for 5-HT and VTX in *Xenopus* oocytes expressing h5-HT3A (n = 5), determined by fitting the current desensitization phase to a monoexponential function (Materials & methods). Representative fits (broken line) of 5-HT (middle panel) and VTX (right panel) desensitization are shown. **E.** Potency of VTX inhibition at h5-HT3A (blue), r5-HT3A (orange) and gp5-HT3A (red) receptors. VTX concentration-inhibition curves were generated using FLIPR Blue membrane potential assay (Materials and methods) to measure responses to 5-HT (30 μM) in the presence of increasing concentrations of pre-applied VTX. Data points represent the mean ± SEM from 5 individual experiments. **F.** Representative traces of h5-HT3A, m5-HT3A r5-HT3A and gp5-HT3A expressed in *Xenopus* oocytes during a 3-step sequential protocol applying twice 10 μM 5-HT and 10 μM VTX in between. **G.** Representative trace from an oocyte expressing h5-HT3A illustrating the persistent inhibition of VTX.

### The m5-HT3AR receptor in complex with VTX adopts an inhibited conformation

In order to grasp the molecular determinants of VTX action, we next sought to obtain a structure of the drug-receptor complex. We initially focused on the mouse receptor. The m5-HT_3A_R was expressed in mammalian cells, solubilized and purified in the detergent C12E9, and imaged by cryo-EM in the presence of 10 μM VTX. A 3D reconstruction was obtained at 3.1 Å resolution (Fig. 2, Fig. SI1-3 and Table SI3) where most of the protein residues are well-resolved, except for the intracellular intrinsically disordered stretch. The structure follows the classical pLGIC architecture, with subunits symmetrically arranged around a central ion pathway. A bound VTX molecule is clearly visible in each of the five orthosteric binding sites. VTX orientation can be assigned unambiguously. No additional densities that could correspond to VTX bound to secondary allosteric sites were observed. The global conformation is largely similar to the conformations previously determined in the presence of -setron antagonist drugs (0.79 Å RMSD for C_α_ atoms with the representative palonosetron-bound structure 6Y1Z, Fig. 2B). This inhibited conformation features a closed pore with a 20-Å long hydrophobic constriction formed by side chains of M2 residues at positions 9’, 13’ 16’ and 17’ (Fig. 2D). To further assess the stability of the m5-HT_3_/VTX complex it was submitted to unrestrained MD simulations for 5×1 μs. The pore is dewetted at the gate level and ion permeation is blocked (Fig. SI4).

**Fig. 2.**
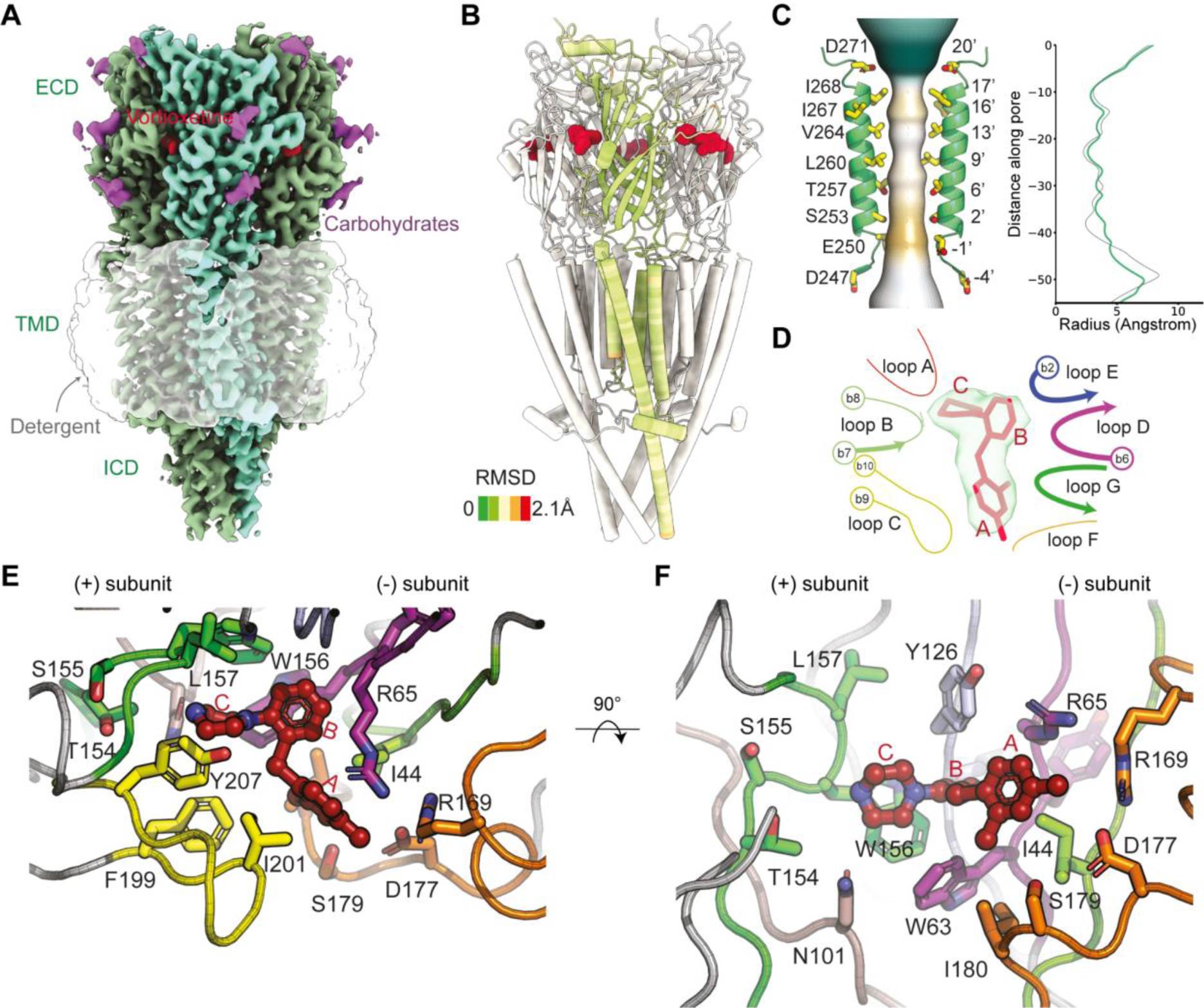
Structure and conformation of the mouse 5-HT3A receptor in complex with VTX. **A.** Cryo-EM and **B.** Cartoon representation of the overall structure of the receptor from the perspective of the membrane. One subunit is colored according to its RMSD with the inhibited palonosetron-bound conformation (after superimposition of the whole pentamers), while the four others are in white. VTX is colored in red. **C.** Close-up view of two opposing pore helices with lining residues depicted as sticks; the surface represents the ion pore and is colored by hydrophobicity. Pore radii are graphed next to the structure (green line, VTX-bound, grey line, palonosetron-bound). **D.** Quality of the Coulombic potential map around VTX. The approximate position of binding elements A to G is schematized. **E-F.** Top and side close-up views of the mouse receptor binding site, with VTX in ball-and-sticks and residues within 4 Å of it depicted as sticks. In the top view, loop E side chains are removed for clarity. In the side view, loop C is removed for clarity. The red A, B, and C labels refer to the VTX cycles.

The two subunits around each neurotransmitter binding site (named the principal and complementary subunit, respectively) both contribute elements to form the binding pocket. These elements are traditionally termed loops A-G (Fig. 2D) even though some are actually portions of β-strands. In the principal subunit, loops A and B line the back of the site while loop C extends as a cap over the bound VTX. In the complementary subunit, β-strands loop E, D and G form the interior of the cavity while loop F lies at its mouth. The piperazine ring of VTX (denoted ring C, Fig. 1A) contains an amine group that penetrates deep into the binding site (Fig. 2E,F). The piperazine occupies a similar location to the positively charged amine group that is found in classical 5-HT_3_ receptor ligands such as the primary amine group of serotonin or the azabicyclo ring a of palonosetron (Fig. SI5). At this location in the binding pocket, the charged nitrogen interacts with the main chain carboxyl of S155 from loop B (3.4 Å distance) and the side chain of N101 from loop A (4.5 Å distance to its tip). In addition, a cation-Pi interaction is observed with the indole group of W156 from loop B (5 Å distance to the center of the aromatic cycle), which is an archetypical hallmark for ligand binding of 5-HT_3_ and nicotinic acetylcholine receptors (Beene et al., 2002; Xiu et al., 2009).

The central phenyl group of VTX (ring B) also penetrates deep into the pocket but is positioned close to residues within loop D, E, and F in the complementary subunit (Fig. 2E,F). Most 5-HT_3_R antagonists do not interact directly with the loops D and E, but instead very often feature H-bond acceptors that form water-mediated interactions (Thompson, 2013). Interestingly, ring B of VTX interacts directly with the backbone of Y64 and is sandwiched between the side chains of W63 and R65 from loop D. In addition, Y126 from loop E appears to make a T-shaped **π-π** interaction with ring B (Fig. 2E, F). The dimethylphenyl group of VTX (ring A) is positioned at the binding pocket entrance, where it remains partly exposed to the solvent. Here, ring A is sandwiched between hydrophobic residues I44, I180, and I201 from loops G, F, and C, respectively.

The m5-HT_3A_R binding pocket residues directly involved in interactions with VTX are summarized in Fig. SI6A. Most of these residues are conserved in the human receptor (the global sequence identity is 85%), suggesting that VTX interacts similarly in h5HT_3A_R. Indeed, the VTX binding mode observed in the m5-HT_3A_R structure is almost identical to a hypothetical binding mode established in a homology model of h5HT_3A_R (Ladefoged et al., 2018). Thus, although the VTX-bound m5-HT_3A_R structure can substantiate that VTX inhibits the mouse receptor via a classical pLGIC antagonist mechanism, the structure does not immediately reveal how the VTX can act as an agonist at the human receptor. Therefore, we set to obtain the structure of h5-HT_3A_R in complex with VTX.

### The Cryo-EM structures of the human receptor feature a distorted transmembrane domain

We imaged the human 5-HT_3A_ receptor stabilized in detergent in the absence of ligand. From this apo dataset, we could isolate three main populations of particles (Fig. 3A and Fig. SI7). A small fraction of the receptors exists as tetramers, with a subunit missing. The reconstruction refined from that fraction has a low resolution. A previous cryo-EM study of the glycine receptor proposed that tetramers could correspond to relevant intermediates along the biosynthesis pathway (Zhu and Gouaux, 2021). However, a more direct interpretation here is that tetramers simply correspond to broken particles where a subunit has been expelled during sample preparation. Approximately another third of the particles yields a reconstruction that possesses the classical features of an apo resting state. The structure built therein possesses a conformation similar to that of the apo or inhibited mouse receptor structures (Cα RMSD of 1.7 and 1.3 Å, respectively, with 6BE1 and 6YZ1), characterized by a closed hydrophobic gate in the TMD, and similar subunit/subunit interface organization. Even though this structure is of interest, it is not further discussed here where the focus will remain on vortioxetine. The third population of receptors yields a reconstruction where the 5-fold symmetry is broken in the TMD while it is mostly preserved in the ECD. The TMD is distorted and adopts a non-physiological conformation where two wide gaps are created at interfaces between a block of 3 subunits and a group of 2 subunits. This distortion is reflected in the buried interface area between subunits. The two gaps correspond to TMD/TMD interfaces that bury surfaces of 0 and 395 Å^2^, while the other three interfaces represent 799, 916, and 974 Å^2^. In line with the symmetry preservation at the ECD levels, all ECD/ECD interfaces are in the same range of 1568±22 Å^2^. The densities at the level of transmembrane helices near the gaps are shallow, again indicative of a distorted flexible structure. Furthermore, the pore lumen is wide (Fig. SI4), with a diameter -to the extent that it can even be trustfully calculated-equivalent to that of the super-open glycine receptor structure (Du et al., 2015). Experimental conditions, combined with an intrinsic fragility of the human receptor, likely distort the TMD. A second defining feature of the structure is the specific organisation of the subunit/subunit interface in the ECD. It is dissimilar to the apo resting conformation but closely resembles the serotonin-bound structures of the m5-HT3AR (see below).

**Fig. 3.**
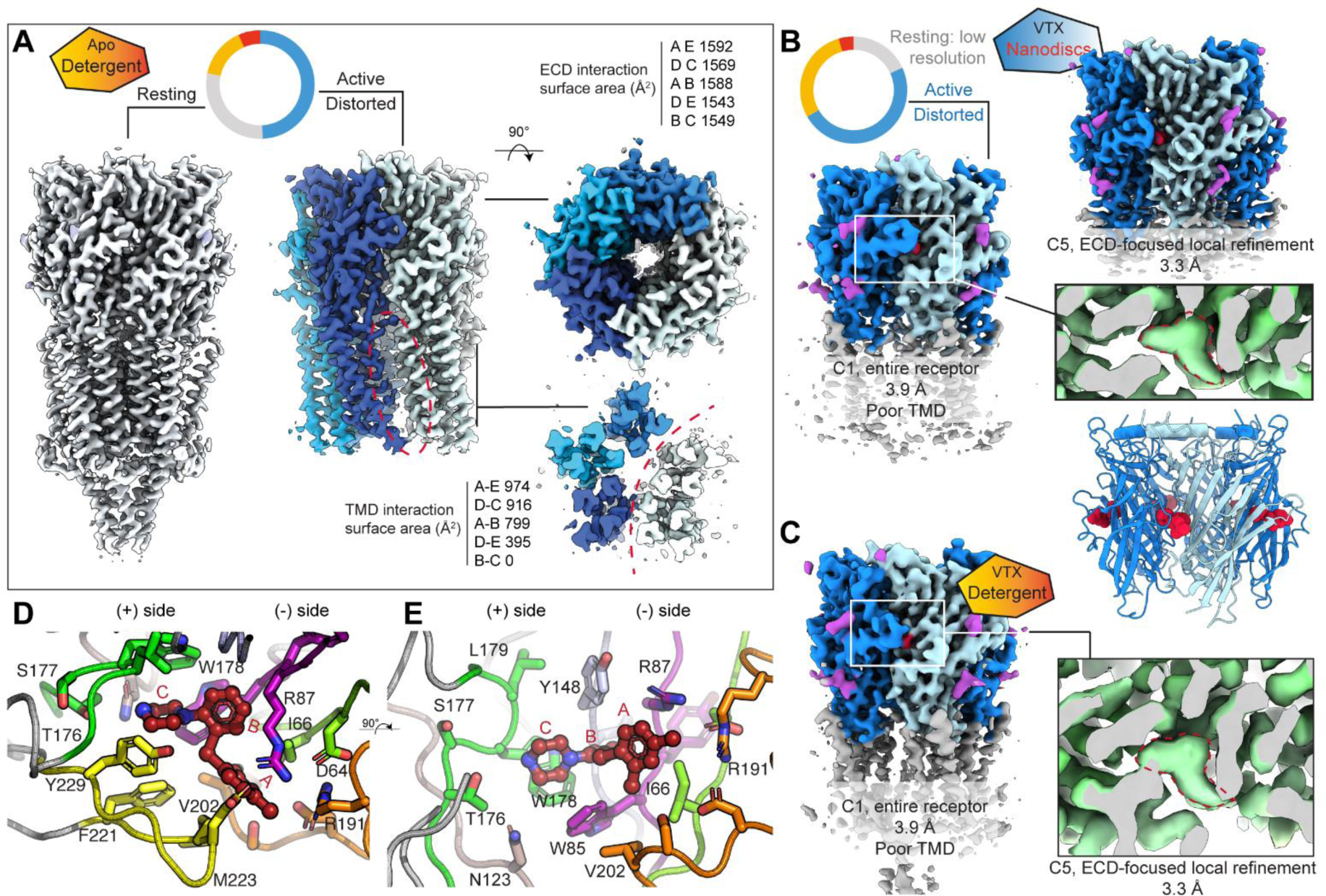
Structures and conformations of the human 5-HT3A receptor. **A**. Cryo-EM maps of the human apo receptor solubilized in detergent. The pie chart indicates the repartition of particles into four categories: junk (red), tetrameric broken receptors (orange), resting state (grey), and a state called active-distorted (blue). The resting state is similar to that of the mouse 5-HT3A receptor, while the active-distorted one presents two wide gaps between subunits in its transmembrane domain (dashed pink lines). At the ECD level, the C5 symmetry is preserved but is broken in the TMD **B.** Cryo-EM maps and structure of the human receptor in complex with VTX, reconstituted in nanodiscs. In that dataset, about half of the particles correspond to the active-distorted state where VTX is bound in the site (inset). The other conformations do not refine to a good resolution. **C.** Cryo-EM map of the human receptor in complex with VTX solubilized in detergent. Only the active-distorted state could be obtained. **D.** Top and **E.** Side close-up views of the human receptor binding site with VTX in ball-and-sticks and residues within 4 Å of it depicted as sticks. In the top view, loop E side chains are removed for clarity. In the side view, loop C is removed for clarity.

Altogether, the apo dataset for the human receptor yielded two reconstructions of good quality: an expected resting closed conformation (similar to what is observed for the mouse receptor) and an unexpected state (denoted active/distorted from here on), with an ECD conformation akin that of the serotonin-bound m5-HT_3A_R structures and with gaps between subunits in the TMD.

Following cryo-EM imaging of detergent-stabilized h5-HT_3A_R in the presence of VTX, one 3D reconstruction was obtained at 3.8 Å resolution for the whole human receptor, which corresponds to the active/distorted conformation (Fig. 3C and Fig. SI8). Densities for VTX are clearly apparent at the interfaces between well-resolved ECDs, but the densities for the transmembrane helices are poor. An ECD-focused local refinement yielded a 3.3 Å resolution reconstruction of this domain (Fig. 3C inset). Given that the detergent-stabilized h5-HT_3A_R could adopt the active/distorted conformation both in the absence and in the presence of VTX, we wondered if replacing the detergent environment with a lipidic one might restore the TMD structure integrity. This approach has previously been applied successfully in cryo-EM studies of human GABA_A_ receptors, in which TMDs were also distorted in detergent but adopted apparently native conformations in nanodiscs (Laverty et al., 2019; Zhu et al., 2018). Therefore, we reconstituted the human receptor in MSP1E3D1 nanodiscs in the presence of soy lipids. Imaging was initially hindered by receptor-containing nanodisc particles adopting preferential orientations in the ice layer (data not shown), which was partly overcome by adding a large high-affinity binder targeting the nanodisc scaffolding protein (termed ‘megabody’) (Uchański et al., 2021). The dataset for the disc-reconstituted receptor in the presence of VTX also featured a population of tetramers and a resting-like state (low resolution). The best reconstruction corresponded to the active/distorted conformation (3.9 Å resolution, Fig. 3B and Fig. SI9). In this reconstruction, the ligand density can be clearly observed in the binding sites, and again the densities for the transmembrane helices are poor. Upon local refinement focused on the ECD, the nominal resolution achieved was 3.3 Å, which allowed the ligand pose to be unambiguously modeled (Fig. 3B inset).

Because of the poor TMD densities, only the ECDs of the human receptors in complex with VTX were built, and only those ECDs are described from here on. At the obtained resolutions, the VTX-bound structures of the h5-HT_3A_R in detergent and in nanodiscs are essentially equivalent at the binding site level. Additionally, the observed binding mode is similar to that previously predicted by the authors (binding mode referred to as C11 in (Ladefoged et al., 2018), which has a heavy atom RMSD of 1.2 Å).

### The human VTX-bound structure possesses an agonist-bound-like conformation in the ECD in line with VTX causing desensitization

In general, pLGICs, including the 5-HT_3A_ receptor, possess some of the hallmark features of allostery. First, the receptor functions by visiting different states populated in a dynamic equilibrium that ligands can shift. Second, the receptor is composed of subunits assembled in a pseudo-symmetric fashion, and a large part of what structurally defines a state lies in the quaternary organization. Subunit/subunit interface rearrangements are major drivers of the transitions between states. In all 5-HT_3_ receptor structures obtained so far, the arrangement of the ECD subunit/subunit interface is a strong marker that differentiates antagonist-bound structures from agonist-bound structures. We, therefore, scrutinized the quaternary organization of VTX-bound receptors. When mouse and human subunit ECDs were superimposed, the backbone RMSD was low for the entire β-stranded core of the subunit used for superimposition (Fig. 4A), indicating that the β-strands move as a rigid body. A few loops did show significantly higher deviations, indicative of a degree of tertiary deformation (see below). Conversely, the backbone RMSD for the neighboring principal subunits featured a strong gradient from ∼0.5 Å on the top of the receptor to ∼2.5 Å at membrane proximal loops (Fig. 4A). Compared to the inhibited m5-HT_3A_R conformation, the h5-HT_3A_R conformation features a rocking motion of one subunit versus the other, associated with a closure of the binding pocket and large motions of loops juxtaposed with the TMD: the Cys-loop, the β1-β2 and β8-β9 loop (Fig. 4B). Next, we overlaid the VTX-bound m5-HT_3A_R and h5-HT_3A_R structures with a range of previously obtained m5-HT_3A_R structures, again using only one ECD for superimposition (Fig. 4C). When looking at the neighboring principal subunit, known antagonist-bound structures clustered with the m5-HT_3A_R structure while known agonist-bound structures clustered with the h5-HT_3A_R structure.

**Fig. 4.**
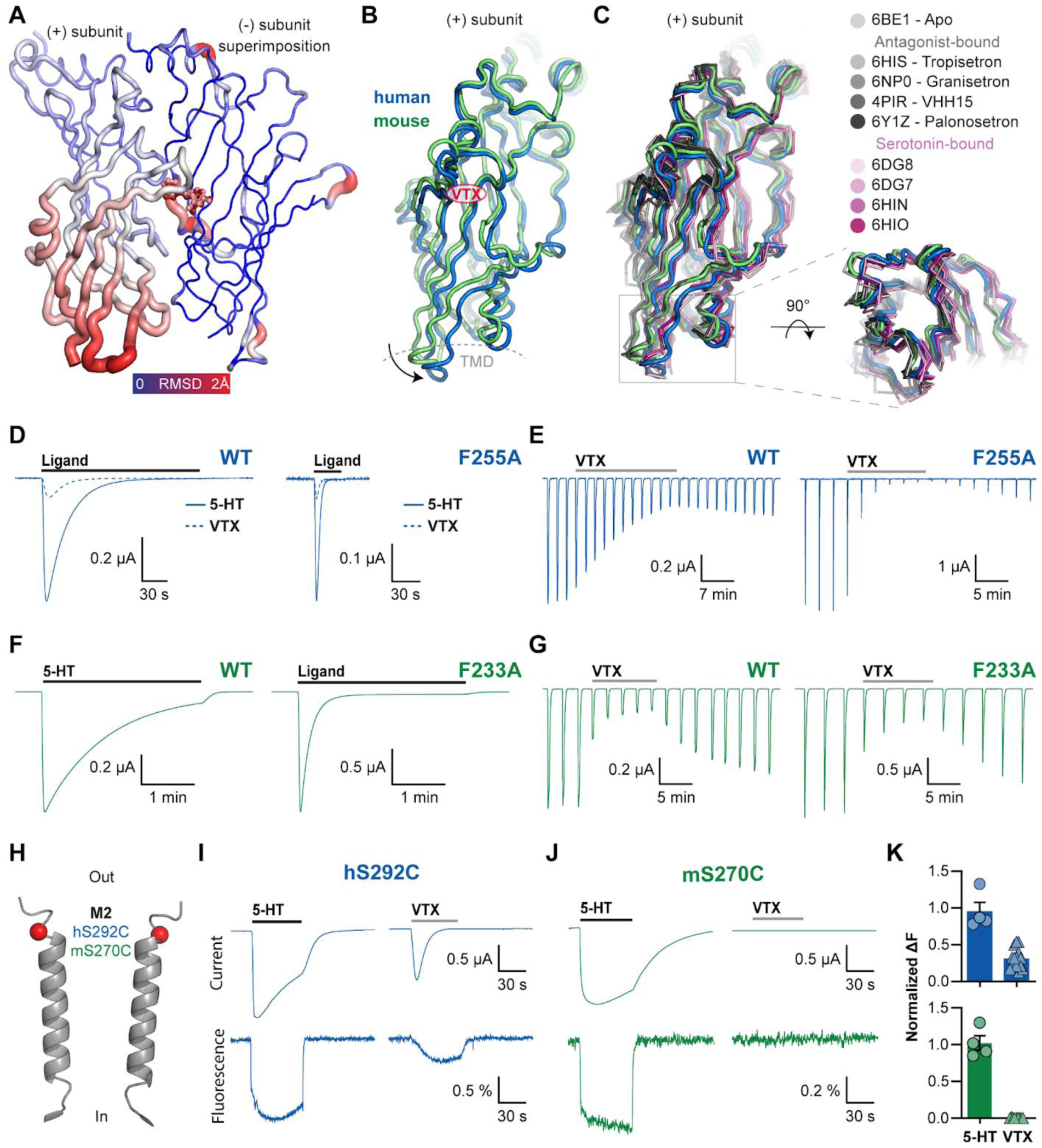
Structural and functional data support the hypothesis of VTX acting through desensitization in h5-HT3AR. **A.** Cartoon representation of two neighboring ECDs, after alignment of the complementary subunit of the human and mouse 5-HT3A receptor. The cartoon is color- and radius-coded by RMSD, illustrating the direction of the quaternary motion. **B.** Overlaid principal subunit ECDs, depicted as green (mouse) and blue (human) cartoons, after alignment of the complementary subunit. Clear deviations are observed at the ECD/TMD interface. **C.** Overlay of the principal subunit ECD representation for all known inhibited conformations (shades of grey) and active agonist-bound ones (shades of pink). Inhibited structures cluster with the mouse VTX-bound structure, while the activated structures cluster with the human VTX-bound structure. **D.** Response elicited by a 3-minutes application of 10 µM serotonin (full line) or 10 µM VTX (dotted line) on oocytes expressing the human wild-type receptor. Response elicited by a 30-second application of 10 µM serotonin (full line) or 10 µM VTX (dotted line) on oocytes expressing the human F255A receptor. The current decay associated with desensitization is accelerated for both ligands. **E.** Progressive inhibition of serotonin-elicited currents (10-second pulses at 10 µM) by a 22-minute (resp. 10-minute) long application of 100 nM VTX on oocytes expressing the human wild-type 5-HT3A receptor (resp. the F255A receptor). The inhibition proceeds faster in the mutant receptor. **F.** Same as D. with the wild-type and the F233A mouse 5-HT3A receptor. Similarly, the mutation strongly accelerates the current decay associated with desensitization. **G.** Same as E with the wild-type and the F233A mouse 5-HT3A receptor. In contrast with the human mutant receptor, the progress of inhibition is not accelerated by the mutation. **H.** Position of the S19’C position in the M2 helix. **I-J.** Conformational changes around S19’C during serotonin (5-HT) and VTX application probed by VCF. Representative simultaneous recordings of current (ΔI) and fluorescence (ΔF) responses to consecutive 100s applications of 5-HT and VTX to oocytes expressing S292C-h5-HT3AR (I) or S270C-m5-HT3AR (J) mutant receptors labeled with MTS-TAMRA. Ligand applications are separated by a 5 minutes wash-out period (not shown). **K.** Summary ΔF response amplitudes from two consecutive applications of 5-HT (circles) or 5-HT followed by VTX (triangles). Each data point represents the response amplitude to the second ligand application normalized to the initial application of 5-HT. Bars represent mean ± S.E.M.

These results support the idea that, at the ECD level, VTX stabilizes an agonist-bound active-like conformation at the human receptor. However, the seemingly artefactual TMD distortion does not allow a definitive conclusion, let alone attribute the h5-HT_3A_R/VTX structure to the desensitized state. Therefore, in an attempt to complement the structural findings, we next used two distinct functional approaches, the results of which both strengthen the hypothesis that VTX stabilizes a desensitized state at h5-HT_3A_R.

First, we identified a TMD mutation that greatly increases the desensitization rate (τ_d_) of the human and mouse receptors. Mutation of a conserved Phe located in the middle of the M1 transmembrane helix to Ala (F255A in h5-HT_3A_R, F233A in m5-HT_3A_R) increases τ_d_ for 5-HT for both receptors (Fig. 4D, F). This increase reflects a shift of the equilibrium between states favoring entry into the desensitized state. We devised a TEVC protocol where short serotonin pulses (10 µM for 10 sec) are applied combined with a several-minute-long VTX application (Fig. 4E, G) that allows tracking of the on-rate of the inhibition of serotonin-induced currents by VTX. The effect of VTX was essentially the same at wild-type and F233A mouse 5-HT_3A_ receptors. This result is in line with the idea that the inhibitory effect of a classical antagonist should not depend on the receptors’ desensitization properties. By contrast, the h5-HT_3A_R F255A mutation significantly altered the effect of VTX compared to that at the wild-type receptor. The on-rate was accelerated, and the degree of apparent steady-state inhibition was higher, in good agreement with VTX stabilizing a desensitized state.

Second, we conducted voltage-clamp fluorometry (VCF) at the h5-HT_3A_ and m5-HT_3A_ receptors utilizing conjugation of the environment-sensitive fluorophore 5-carboxytetramethylrhodamine methanethiosulfonate (TAMRA) to Cys residues inserted at a previously established reporting position located on the top of the M2 helix (S19’C or S270C-m5-HT_3A_R and S292C-h5-HT_3A_R, respectively) (Polovinkin et al., ^2^018). At this position, TAMRA emission is sensitive to the conformational changes in the ion pore and decreases during channel entry into the active-open and desensitized-closed states. Following TAMRA labeling, we recorded S19’C receptors’ current and fluorescence change (ΔF) to the consecutive application of saturating concentrations of serotonin (100 µM) and VTX (10 µM). At the human receptor, both 5-HT and VTX application caused decreases in fluorescence during current activation to a steady-state level that remained unchanged during the current desensitization phase (Fig. 4h). The fluorescence change during the VTX application was consistently 3-fold smaller than the serotonin-evoked change (69 ± 5 %, n = 9), in line with VTX being a partial agonist. This result strongly suggests that prolonged applications of 5-HT and VTX induce similar structural rearrangements around S19’C of the M2 helix and, thus, that VTX stabilizes a desensitized state. In contrast, VTX did not change fluorescence at the mouse receptor, as expected for a classical antagonist that cannot communicate allosterically to the gating region.

In conclusion, we provide three converging lines of evidence (four structures, electrophysiology on fast-desensitizing mutants, VCF of an M2-attached probe) that collectively support the hypothesis of VTX stabilizing a desensitized state similar to the state stabilized by the natural agonist serotonin, at the human receptor.

### Loops C and F contain determinants of VTX activity

Following the analysis of the global receptor conformations, we next focused on dissecting the structural determinants in the human and rodent receptors for the differential VTX activity. As previously mentioned, a comparison of the VTX-bound h5-HT_3A_R and m5-HT_3A_R structures shows that the binding mode of VTX is largely similar in the two binding pockets (Fig. 5A-B). Ring A and the central ring B are orientated almost completely identically with atom-position RMDS values in the 0.5-1 Å range, whereas ring C is slightly offset (atom-position RMSDs in the 1.5-2 Å range) as the dimethyl phenyl is positioned closer towards loop E in m5-HT_3A_R (Fig. 5A-B). In addition to similar orientation, ring A-C forms near-identical ligand/protein interactions in both receptors as they are surrounded by residues highly conserved between h5-HT_3A_R and m5-HT_3A_R (Fig. 5A and SI11B). Specifically, the piperazine ring C is surrounded by identical polar residues (N123/N101 on loop B, W178/W156 on loop E, and W85/W63 on loop D, in h5-HT_3A_R and m5-HT_3A_R, respectively). The central ring B is sandwiched between two identical aromatic residues (W85/W63 on loop D and Y148/Y126 on loop E), and, at the pocket entrance, ring A is surrounded by two similar hydrophobic residues on loop F (V202/I180) and loop C(M223/I201), in addition to conserved Ile (I66/I44) residues on loop G. Also, the pocket conformations are similar to the tertiary positions of the loop backbones and contact residue side chains superimposing almost entirely, with the most pronounced difference being loop C which adopts a more capped conformation in the human receptor, which makes the human pocket slightly more closed (Fig. 5A-B).

**Fig. 5.**
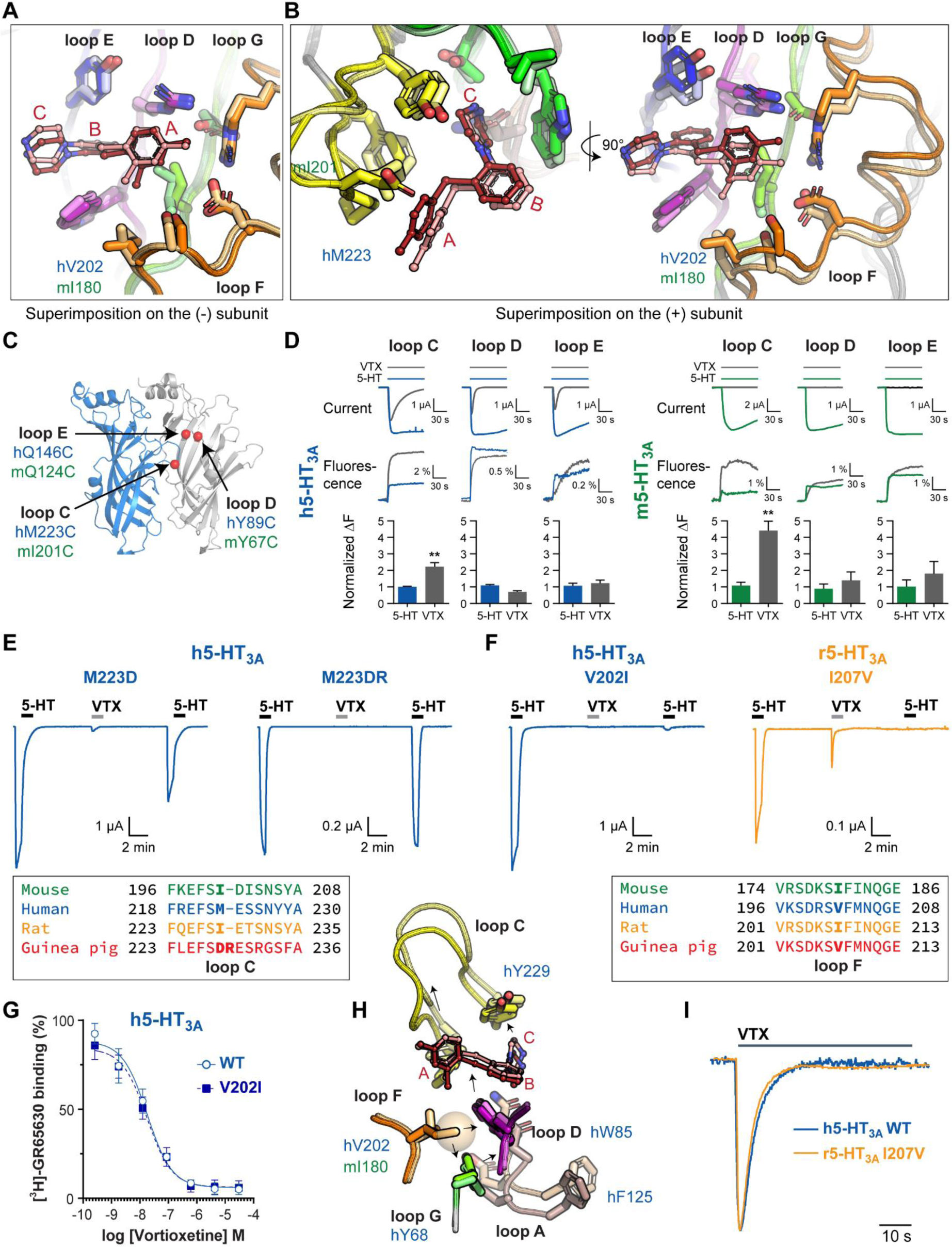
Comparison of VTX binding at the mouse and human 5-HT3A receptors. **A.** Overlay of the mouse (pale colors) and human (vivid colors) receptor structures depicted as cartoon and sticks for the first shell of interacting residues, when complementary subunits are superimposed. **B.** Overlay when principal subunits are superimposed. **C.** Location of VCF reporter positions around the orthosteric binding pocket. **D.** Conformational changes in the binding site probed by VCF. Representative simultaneous recordings of current (ΔI) and fluorescence (ΔF) responses to applications of 5-HT and VTX to oocytes expressing h5-HT3AR mutant receptors labeled with MTS-TAMRA. The bar graphs represent the mean ± S.E.M ΔF response amplitudes. **E.** Representative traces of h5-HT3A mutant receptor harboring guinea pig residues at the tip of loop C during a 3-step protocol applying twice 10 μM 5-HT and 10 μM VTX in between. (equivalent traces for the wild-type receptor are depicted in Fig. 1F). The inset depicts the local sequence alignment. **F.** Traces for the loop F mutants that revert the VTX agonist phenotype, hV202I, and rI207V receptors. **G**. Representative concentration-inhibition curve for VTX displacement of [3H]-GR65630 binding at membranes expressing WT h5-HT3A and V202I receptors. Error bars indicate the SD for a single-day experiment conducted in quadruplicate. **H.** Speculative mechanism explaining the V202I mutant phenotype. Starting from the added methyl group (brown sphere) the arrows represent the chain of conformational deviations that reposition loops A and C and ultimately prevent activation. **I.** Overlay of normalized VTX responses in WT h5-HT3A (blue) and r5-HT3A-I207V receptors (orange) to illustrate the similar desensitization time course.

The comparisons of the h5-HT_3A_R and m5-HT_3A_R structures do not reveal major differences in how VTX interacts with the receptors that explain its differential action. Therefore, to explore VTX interactions with the binding pockets in a dynamic environment, we subjected the VTX-bound h5-HT_3A_R and m5-HT_3A_R structures to five sets of 1 μs MD simulations (Fig. SI4). These simulations did reveal two major differences in the behavior of the ligand at the two receptors. First, we observed that stabilizing hydrogen bond (HB) interactions to the charged amine in ring C appeared to form differently between mouse and human receptors, with HBs forming mostly with N123 in m5-HT_3A_R and with S177 in h5-HT_3A_R (Fig. SI4E). Second, we observed a consistent difference in the stability of loop C between the mouse and human simulations. Specifically, the distance of the center-of-mass of loop C to loop D on the complementary subunit appeared to fluctuate more in m5-HT_3A_R than in h5-HT_3A_R over the simulation periods (Fig. SI4B). Notably, this distance-distribution difference relates to a difference in the distribution of dihedral angles assumed by the sulfide bond (C-S-C) between rings A and B in the simulations. Specifically, in all the simulations, VTX assumed conformations with mainly two highly populated angles, one at approximately −40° corresponding to an “upwards” orientation of ring A and one at approximately 40° corresponding to a “downwards” orientation that brings ring A closer to loop F (Fig. SI4F). During the simulations, ring B remains stable in both m5-HT_3A_R and h5-HT_3A_R, locked by the identical interaction network. Therefore, the downward/upward dihedral angle configurations dictate the positioning of ring A. The downwards conformation dominates in h5-HT_3A_R during parts of our simulations; however, not enough to be statistically significant on the current simulation timescale. Nonetheless, an important consequence of the downward and upwards orientation seems to be how ring A interacts with residues on loops C and F. Interestingly, of all 54 residues within 4 Å distance of VTX, the V202/I180 position is among the three positions that are non-conserved among human and mouse 5-HT_3A_Rs (Fig. SI6B). The Ile and Val side chains only differ by a single methyl group, yet, this slight difference may be a determinant for the differential orientation of ring A between m5-HT_3A_R and h5-HT_3A_R and perhaps underlie the functional differences of VTX.

Although the cryo-EM structures show VTX to interact similarly with m5-HT_3A_R and h5-HT_3A_R, the MD simulations suggest that VTX may influence the conformational behavior of the binding region differentially during steady-state binding conditions. To test this idea, we used VCF to probe if VTX binding elicits local conformational changes around the pocket differentially in mouse and human 5-HT_3A_R in a live-cell membrane environment. Specifically, in human and mouse receptors, we labeled Cys residues inserted into loop C (M223/I201), loop D (Y89/Y67), and loop E (Q146/Q124) with TAMRA, which previously have been shown to report conformational changes in the binding pocket region in h5-HT_3A_R, as well as other pLGICs (Chang and Weiss, 2002; Lefebvre et al., 2021; Munro et al., 2019; Muroi et al., 2006; Pless and Lynch, 2009). From all three reporter positions in h5-HT_3A_R, sequential application of saturating concentrations of 5-HT and VTX elicited increases in fluorescence that were similar in direction and amplitude, except for a small but significant increased VTX response amplitude for the loop C position (Fig. 5D). However, the VCF analysis overall indicates that VTX and 5-HT stabilize largely similar pocket conformations. For m5-HT_3A_R, a similar picture was observed for the loop D and E reporter positions with VTX and 5-HT evoking similar ΔF responses (Fig. 5D). However, for the loop C reporter position, the magnitudes of the ΔF for VTX and 5-HT were substantially different with VTX producing approximately 5-fold larger increases in TAMRA fluorescence (Fig. 5D). These data support that VTX and 5-HT stabilizes the binding pocket in different conformations in the mouse receptor, in line with their different functional effects.

The loops A–C in the principal subunit and D–F in the complementary subunit are highly conserved among the human and the rodent 5-HT_3A_Rs (Fig. SI6B). We tested if the nonconserved residues determine differential VTX activity by switching non-conserved residues from one species to another and assessing the effect on VTX phenotype, focusing first on human and guinea pig receptors, where VTX activity differs most in terms of agonist activity and potency. The h5-HT_3A_R and gp5-HT_3A_R differ at eight positions in loops C, F, and G (Fig. SI6B). We first mutated all of these in h5-HT_3A_R to become identical to gp5-HT_3A_R and determined the VTX response phenotype of the mutant receptor (h5-HT_3A_R-gp1, Fig. SI6C). Interestingly, h5-HT_3A_R-gp1 showed a VTX phenotype near identical to WT gp5-HT_3A_R with no VTX agonist response, a 40-fold increase in IC_50_, and near-complete recovery of 5-HT response from VTX inhibition (Fig. SI6C). Five of the eight differing residues are constrained to a 7-aa stretch in loop C with the sequence MESSNY in h5-HT_3A_R such that the N-terminal M (M223) is replaced by a DR motif, extending loop C by a single residue, and the C-terminal SNY motif is replaced by RGS (Fig. SI6C). Among these positions, M223 is the only residue within 4Å distance of VTX (Fig. SI6A-B). Therefore, we focused on the M/DR difference and replaced M223 in h5-HT_3A_R with aspartate. The resulting h5-HT_3A_R-M223D mutant also showed a guinea-pig-like VTX phenotype with no agonist response, partial recovery from VTX inhibition, and ∼20-fold increase in VTX IC_50_ (Fig. 5, Fig. SI6C, and Table SI3). Further insertion of the Arg residue created the h5-HT_3A_R-M223DR mutant, which added full recovery from VTX inhibition, thus completely reverting the human receptor to a guinea pig VTX phenotype (Fig. 5, Fig. S11C, and Table SI3) and identifying the M/DR difference in loop C as a sole determinant for VTX activity differences.

We then focused on the determinants for the differences in human versus rat/mouse phenotype. Overall, the binding elements in m5-HT_3A_R and r5-HT_3A_R are identical, except for a single position in loop D and two consecutive positions in loop C (Fig. SI6B). Specifically, in m5-HT_3A_R loop D contains a Tyr at position 67 (Y67), which is Phe in r5-HT_3A_R (F94), and in loop C, an Asp (D202) and an Ile (I203), which in r5-HT_3A_R are Glu (E229) and Thr (T230), respectively (Fig. SI6B). However, in the m5-HT_3A_R structure, the side chains of these residues point away from VTX or are positioned more than 10 Å from the ligand. Thus, the mouse and rat VTX binding sites can overall be considered conserved, which can explain the near-identical VTX phenotypes of these receptors. Seven residues in the human receptor loops C, F, and G differ from the mouse/rat sequence (Fig. SI6B). Similar to the human/guinea pig mutational analysis, we mutated all of these in h5-HT_3A_R to become identical with r5-HT_3A_R. Indeed, the resulting mutant (h5-HT_3A_R-r1) showed a VTX phenotype nearly identical to WT r5-HT_3A_R, suggesting that one or more of the non-conserved residues are determinants for VTX activity (Fig. SI6d and Table SI1). Among the seven differing residues, only V202 in loop F interacts directly with VTX in the h5-HT_3A_R structure (Fig. 5 and Fig. SI11B), and we, therefore, mutated V202 to the Ile contained in mouse/rat. This mutant (h5-HT_3A_R-V202I) displayed a phenotype nearly identical to WT r5-HT_3A_R (Fig. 5F and Table SI1). Importantly, the IC_50_ for VTX as well as the dissociation constant (*K*_d_) for the radioligand [^3^H]-GR65630 was not changed by the V202I mutation (Fig. 5E and SI12 and Table SI1), showing that the V202I mutation does not abolish the current response to VTX by perturbing ligand binding (Fig. 5F, Fig. SI12 and Table SI1). Individual mutation of the remaining positions in the loop F and in loop C that differ between human and mouse/rat, but are not in contact with VTX, to the mouse/rat identity in h5-HT_3A_R did not change the VTX response phenotype (R200K M204I, M223I, S225T, and Y228S in h5-HT_3A_R; Fig. SI6B and Table SI1
). We further tested the determinant role of the Val/Ile difference by making the reverse I-to-V mutation in the rat receptor. Indeed, the mutant receptor (r5-HT_3A_R-I207V) now displayed rapidly desensitizing agonist responses to VTX (Fig. 5F), with a time course similar to that of the wild-type human receptor (Fig. 5I). Together, these results identify the Val/Ile difference in loop F as the sole determinant for VTX agonist activity at human and rat/mouse receptors.

### Structure-activity relationship of VTX analogues

The species-scanning mutagenesis identified a single Ile/Val difference on loop F as the sole determinant for agonist activity between human and rat receptors. Intriguingly, the V202I mutation only adds a single methyl (-CH_3_) group to the binding pocket yet completely abolishes the current response to VTX. In the VTX-bound m5-HT_3A_R and h5-HT_3A_R structures, this position is the closest to ring A in VTX (Fig. 5A). Therefore, to probe the role of ring A/loop F interaction for VTX agonism, we characterized a series of compounds that share an identical three-ring scaffold with VTX but have different substitutions in positions R_1_, R_2_, and R_3_ in ring A (Bang-Andersen et al., 2011)h5-HT_3A_R and the rodent-like h5-HT_3A_R-V202I mutant receptor, using the membrane potential assay (Fig. SI10). In VTX, ring A has methyl (-CH_3_) groups at the R_1_ and R_3_ positions and the R_2_ position is unsubstituted (Fig. 6A). We first focused on an analog series where -CH_3_ at the R_1_ and R_3_ positions either remained as a substituent or removed or mono- or di-substituted with larger, but still hydrophobic methoxy (-OCH_3_) groups (Fig. SI10A). The resulting structure-activity relationship (SAR) provided two major insights. First, all analogs were found to retain agonist activity at h5-HT_3A_R (Fig. SI10A). This rather large tolerance for substitutions on ring A indicates that direct interaction with functional groups on ring A is not per se determinant for h5-HT_3A_R agonism, which also fits well with the VTX-bound h5-HT_3A_R structure showing the R_1_-R_3_ positions to point out of the binding site (Fig. 6B). Second, at the h5-HT_3A_R-V202I mutant, several analogs rescued agonist activity (Fig. SI10). Specifically, compounds 4j, 4g, 5g, and 5e evoked responses at the V202I mutant similar in magnitude to those of VTX at the WT receptor (Fig. SI10). These analogs all have the R_3_ position unsubstituted, while the R_1_ position remains substituted with either -CH_3_ or -OCH_3_. The R_3_ substituent is closest to the 202 side chain (Fig. 6C,D). This SAR data, therefore, indicate that the lack of VTX agonism in mouse/rat receptors is determined by the additional -CH_3_ in the Val side chain clashing with the R_3_ substituent. This hypothesis is best illustrated by compound 4j, which is identical to VTX except for the single substitution of the -CH_3_ group with hydrogen in R_3_ that rescues agonist response at the h5-HT_3A_R-V202I mutant receptor.

**Fig. 6.**
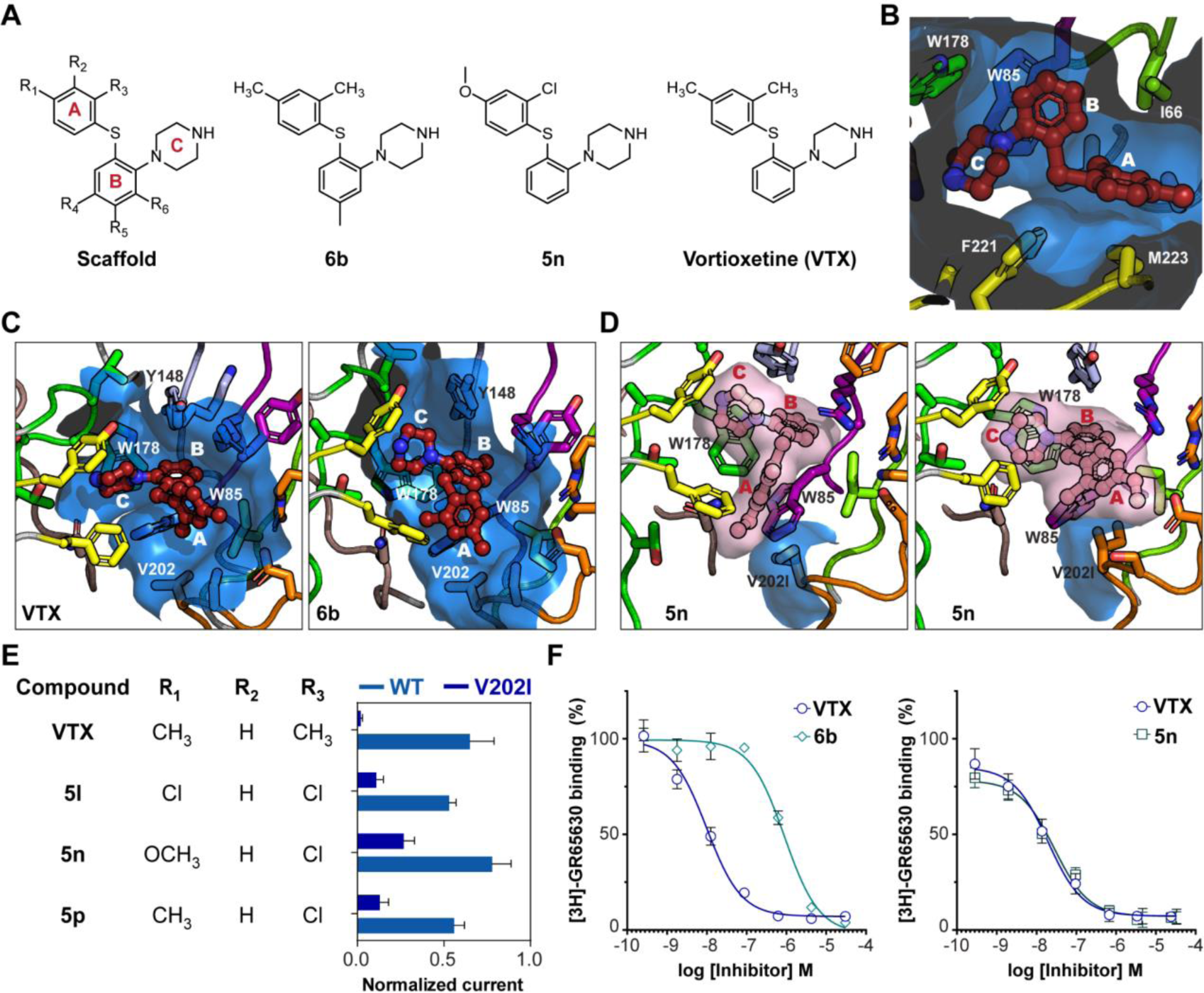
Structure-activity relationship of VTX analogues. **A.** Chemical structures of the vortioxetine scaffold, vortioxetine, 6b and 5n. **B.** Top view of the binding pocket with VTX bound. **C.** Similar side views of the binding pocket with bound VTX and docked 6b compound. **D.** Side views of the binding pocket in the V202I mutant with two docked poses of compound 5n, both of them showing interaction of ring A with I202. **E.** Amplitude of agonist response of 10 μM vortioxetine and analogous compounds, relative to 10 μM 5-HT measured by TEVC. **F.** Representative concentration-inhibition curve for the displacement of [^3^H]-GR65630 by vortioxetine, 6b and 5n at h5-HT3A receptors. Data points represent the mean + SD for a single-day experiment conducted in quadruplicate.

We also screened a second series of analogs with permutations of the substituents in both ring A and B (Fig. SI10). VTX itself does not contain any functional groups on ring B. The resulting SAR data provided two major insights, First, most analogs with a substituent in ring B displayed decreased or complete loss of agonist activity. Adding a -CH_3_ substituent at R_4_ or R_6_ (6a and 6c) lowers agonist response, and a -CH_3_ group at R_5_ entirely abolishes agonism regardless of other substituents (4a, 4b, 4c, 4d, 4e, and 6b). This effect is most striking for analog 6b, which is identical to VTX except for an additional -CH_3_ on R_5_. Radioligand-displacement experiments at h5-HT_3A_R using [^3^H]-GR65630 with 6b showed a 37-fold decrease in *K*_i_ value compared to VTX (*K*_i_ = 24 [13; 44] nM for VTX, *K*_i_ = 880 [290; 2600] nM for compound 6b), indicating that 6b has lower binding affinity than VTX and, therefore, at the 10 µM screening concentration might not saturate the receptor sufficiently for activation. However, increasing the concentration of 6b to 100 µM, thus more than 100-fold higher than the *K*_i_, still did not evoke any response (data not shown). These results clearly show that agonism at h5-HT_3A_R is incompatible with added bulk on ring B. In the h5-HT_3A_R structure, ring B is lined against the backbone of Y86 in loop D in a position optimal for **π-π** and cation-**π** interaction with W85 and R87, respectively (Fig. 3D, E). We docked 6b in the h5-HT_3A_R structure and found the additional -CH_3_ at R_5_ indeed disrupts this interaction network and moves ring B away from Y64 and towards the principal subunit resulting in steric clashes (Fig. 6C). Overall, these findings suggest that VTX agonism requires ring B to assume a very specific orientation around Y86 in loop D.

A second insight came from the observation that several analogs (5n, 5p, and 5l) with chlorine (-Cl) in the R_3_ position rescued agonist response at the V202I mutant (Fig. SI9B), which was confirmed using TEVC electrophysiology (Fig. 6E). We selected compound 5n with the highest activity for further analysis. In addition to having -Cl at R_3_, 5n differs from VTX by having -OCH_3_ in R_1_ (Fig. 6E). Radioligand-displacement experiments showed 5n to have binding affinity at the V202I mutant identical to VTX at WT h5-HT_3A_R (Fig. 6E). Docking of 5n into the V202I mutated binding pocket showed that in contrast to VTX, the R_1_ and R_3_ substituents could interact directly with the Ile side chain. At the same time, rings A and B are positioned identically to VTX in WT h5-HT_3A_R (Fig. 6D). Specifically, in the highest scoring binding mode, the R_1_ and R_3_ substituents almost “wrap” around the Ile side chain, bringing ring A very close to loop F. Conversely, when docking vortioxetine into the V202I mutant receptor, VTX was observed to bind with much more variation within the binding site compared to WT receptors and with no direct interaction with V202I. Only one cluster is similar to the binding mode observed on h5-HT_3A_R with ring B and C in highly similar positions in the binding site as when binding in the WT h5-HT_3A_R; however, phenyl A is consistently pointing upwards in the binding site, i.e., away from V202I.

Taken together, our SAR analysis of activity at WT vs h5-HT_3A_R-V202I mutant further corroborates that the relative conformation and positioning of ring A and B in the pocket are key determinants for agonism or antagonism of this molecular scaffold. Specifically, we hypothesize that the rodent antagonism of VTX is caused by the larger Ile side chain in loop F of the rodent receptors to induce both rings A and B of the ligand to adopt a conformation that prevent the pocket structure from transitioning into the condensed conformation that is necessary to trigger conformational transitions further downstream in the receptor structure towards channel activation.

### Recovery from VTX inhibition

The long-lasting inhibition by VTX has been reported at several instances since the compound discovery. For instance, Bang-Andersen and colleagues initially found that VTX at high concentration (30 µM applied for 30 sec) induces entry in a non-recovering state in human and rat 5-HT_3A_ receptors (Bang-Andersen et al., 2011). The absence of recovery was replicated by Dale and colleagues, using longer applications of a lower concentration of the drug (100 nM for 30 min) on human 5-HT_3A_ receptors, which inhibited serotonin-induced currents for more than two hours in stark contrast with the behavior upon equivalent application of the classical antagonist ondansetron (Dale et al., 2018). The authors hypothesized that there were *“differences in the manner with which ondansetron and VTX bind to the 5-HT3 receptor at the molecular level”*. While our combined structural and functional approach brought insight on the molecular mechanism of VTX at the orthosteric site, the long-lasting inhibition remained unaccounted for. Indeed, VTX binds to the same site as ondansetron, palonosetron, and granisetron, which also possess extremely high affinities for the receptor, but do not irreversibly inhibit activation. We investigated two scenarios that could explain the persistent loss of response following the VTX application. We first asked whether VTX induced receptor internalization as has been reported for some 5-HT_3_ receptor ligands (Ilegems et al., 2004) by fusing h5-HT_3A_R with β-lactamase enzyme (β-lac-h5-HT_3A_), which allowed the measurement of receptor surface levels over time by use of a colorimetric assay (Lam et al., 2013). Exposure to 10 µM VTX for 30 minutes did not reduce the surface expression of βlac-h5-HT_3A_ in both HEK-293 cells and *Xenopus* oocytes (Fig. SI11); thus, disproving the internalization hypothesis.

VTX has an octanol/water partition coefficient (log P_o/w_) of 4.94, and is more lipophilic than granisetron (log P_o/w_=2.6) or palonosetron (log P_o/w_=2.7). We therefore asked whether VTX partition into the oocyte membrane, and thus delayed desorption, could explain the absence of recovery. Initial experiments were conducted using M-ß-cyclodextrin, a compound classically employed for cholesterol depletion in electrophysiology. Because hydroxypropyl-ß-cyclodextrin was also used as a vehicle during preclinical studies for VTX delivery (Guilloux et al., 2013), we reasoned it may deplete an eventual accumulation of VTX in the membrane. Indeed, strikingly faster recoveries were occasionally observed following washes with 20 mM M-ß-cyclodextrin (data not shown), but the effect was too variable to be conclusive. A putative reason for the variability is that M-ß-cyclodextrin influences the receptor function independently of VTX. We therefore designed an experiment comparing the effect of short or long VTX applications. If VTX accumulates into the lipid bilayer, longer applications should yield larger inhibitions; if VTX does not partition in the membrane, then its effect should not depend on the length of application (as long as the rate of drug binding is not a limiting step, which is unlikely given the affinity). During repeated short applications of serotonin (10-second pulses at 10 µM), the effects of 30-second or 5-minute-long application of 100 nM VTX were compared for the m5-HT_3A_ receptor (Fig. 7A, B). The recovery from VTX application showed a biphasic time course in both experiments, however, a clear difference in the proportion of the kinetic components was observed. Specifically, after the short application, a steep initial recovery indicated that fast and slow components were of similar amplitude, whereas the slow component dominated the recovery after the long application. It was necessary to use the rapidly desensitizing F255A construct to perform equivalent experiments on the human receptor. With the WT receptor, we either observed little inhibition at a low concentration (e.g., 100 nM for 30 sec) or the recovery was too slow at higher concentrations (e.g., 1 µM for 30 sec). This behavior stems from the fact that the dose-inhibition curve of the WT h5-HT_3A_R is very steep compared to that of the m5-HT_3A_R. In the mutant F255A h5-HT_3A_R, as already observed for the m5-HT_3A_R, recovery was slowed down after a long application of 9 minutes compared to a short application of 30 sec (Fig. 7C, D). Together, these results suggest that the persistent inhibition of 5-HT_3A_ receptors by VTX is due to the ligand binding to the orthosteric site and stabilizing either an inhibited (in the mouse or the rat receptor) or a desensitized state (in the human receptor) from which the recovery rate is extremely slow due to 1) slow dissociation and 2) sustained release of the drug from the membrane where it tends to partition.

**Fig. 7.**
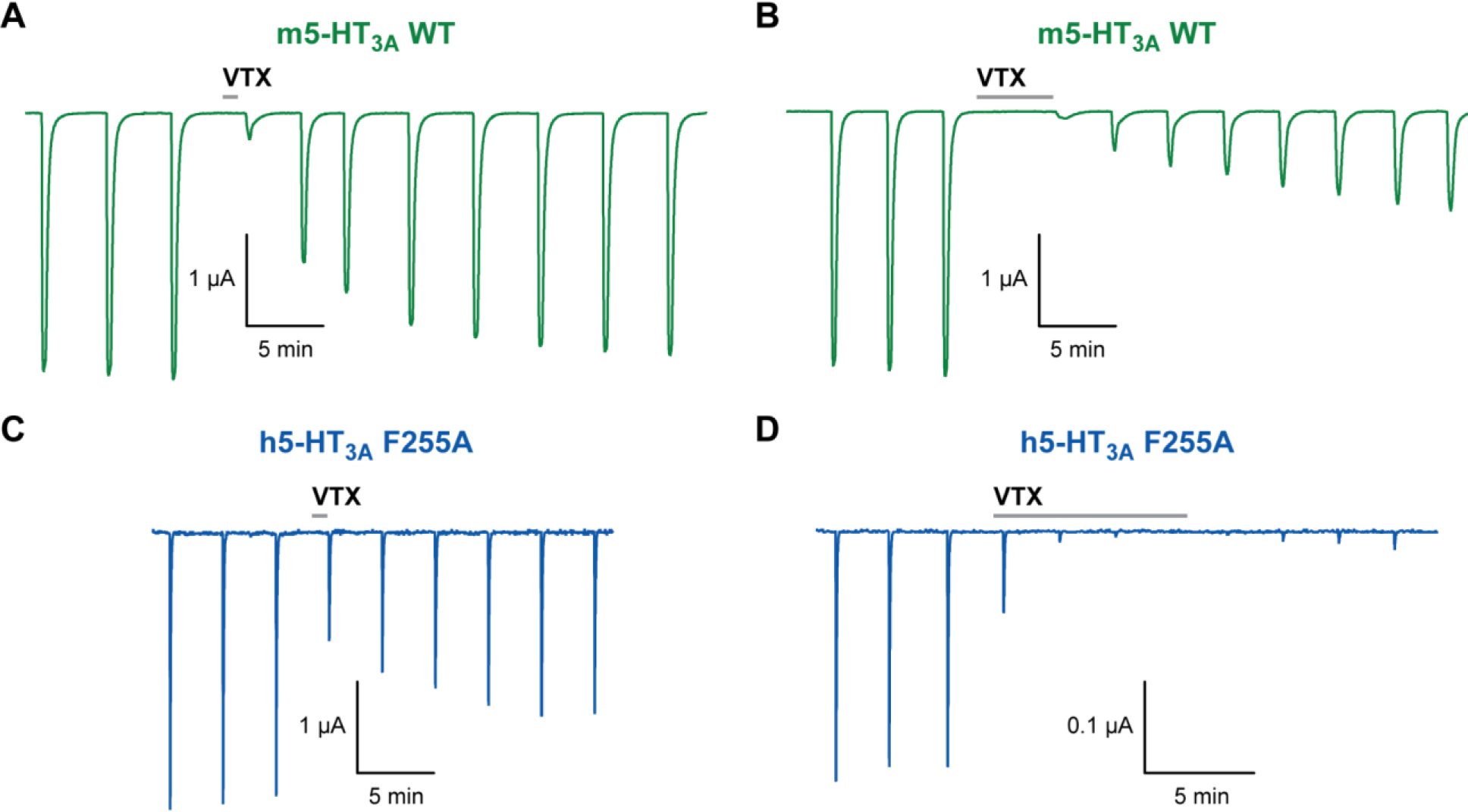
The washout time course of VTX depends on the length of its application. **A.** Recovery of serotonin-elicited currents (10-second pulses of 10 µM) after a short 30-sec application of 100 nM VTX on oocytes expressing the mouse wild-type receptor. The time course of recovery shows a slow and a fast component of equivalent amplitudes. **B.** Recovery of serotonin-elicited currents (10-second pulses at 10 µM) after a longer 5-minute application of 100 nM VTX. The fast recovery component is now dominated by the slow component. **C and D.** Same protocol at human F255A mutant receptors with a short 30 sec and a long 9 min VTX application, respectively. Similar to the mouse receptor, the recovery time course strongly depends on the length of application of VTX.

## Conclusion

VTX stands out in the pharmacology of 5-HT_3_ receptors for three main reasons. First, it is the only prescription drug targeting this receptor that does not treat nausea and emesis but possesses antidepressant properties. Second, it is, to our knowledge, the sole compound displaying differential activities at human and rodent 5-HT_3_ receptors. Third, the absence of, or the very slow recovery from VTX inhibition puts it apart from the equivalently potent antagonists of the setron family.

Using a combination of structural and functional approaches, we have established a molecular basis that explains, for a large part, how VTX operates. In particular, structures revealed a largely common binding mode of the drug at the mouse and human receptors, which, nevertheless, stabilizes two different quaternary states of the ECD. We assigned the mouse receptor structure to the inhibited state without ambiguity, proving that the compound constitutes a competitive antagonist. For the human receptor, we presented a set of converging experiments that support the idea that VTX stabilizes a desensitized state. We could pinpoint residues of loop C and loop F where mutations crucially impact the agonist activity and steady-state inhibition of VTX. Exploring the structure-activity relationship of VTX, we described how substitutions of chemical groups impact VTX activity. We also provided an explanation for the slow recovery of the vortioxetine inhibition, which seems to be caused by the partition of the drug in the membrane in combination with nanomolar affinity. Along the way, we also observed an intrinsic fragility of the human receptor transmembrane domain that led to artefactual TMD conformations in detergent and lipid nanodiscs. The human receptor structures shed some light on how the transmembrane domain of a pLGIC can be unstable and how its structure can deviate from the 5-fold symmetry.

Beyond a mere deep dive into how VTX operates, our findings provide new insight into the molecular basis of orthosteric ligand activity at 5-HT_3_ receptors and the determinants for agonist versus antagonist activity, which may be used in the rational design and development of novel therapeutic agents. We believe there is more work to be done, on the medicinal chemistry side so that 5-HT3 receptors fulfill their potential in neurological disorders (Fakhfouri et al., 2019; Zajdel et al., 2021).

## Acknowledgements

This work was supported by the Danish Council of Independent Research for Medical Sciences (grant number DFF-404-00309), the Lundbeck Foundation (grant number 2017-1655 and 2012-12453), the Carlsberg Foundation and the ERC Starting grant 637733 Pentabrain. U.L-S. was supported by the *Fondation pour la Recherche Médicale* (Grant number SPF201809007073). Computations were made possible through allocations at the Centre for Scientific Computing, Arhus (SCS-Aa). We acknowledge access to the ESRF CM01 Krios microscope. It used the platforms of the Grenoble Instruct-ERIC center (ISBG; UMS 3518 CNRS-CEA-UGA-EMBL) within the Grenoble Partnership for Structural Biology (PSB), supported by FRISBI (ANR-10-INBS-05-02) and GRAL, financed within the University Grenoble Alpes graduate school (Ecoles Universitaires de Recherche) CBH-EUR-GS (ANR-17-EURE-0003). The electron microscopy facility is supported by the Rhône-Alpes Region, the FRM, the FEDER and the GIS-IBISA. We thank Lefteris Zarkadas for his help with cryoEM. F.D. acknowledge the State-Region Plan “Technological Innovations, Modeling and Personalized Medical Support” (IT2MP) and the European Regional Development Funds (ERDF) for generous support.

## Author Contributions

Participated in research design: all authors Conducted experiments: López-Sánchez, Munro, Ladefoged, Pedersen, Colding-Bruun, Lyngby, Schoehn, Neyton, Dehez, Chipot, Nury, Kristensen Contributed new reagents or analytic tools: Bang-Andersen. Performed data analysis: López-Sánchez, Munro, Ladefoged, Pedersen, Colding-Bruun, Lyngby, Neyton, Dehez, Nury, Kristensen Wrote or contributed to the writing of the manuscript: López-Sánchez, Munro, Ladefoged, Nury, Kristensen.

## Materials and Methods

### Materials

Chemicals were from Sigma (St. Louis, MO) except otherwise stated. Dulbecco’s modified Eagle’s medium (DMEM), fetal bovine serum, trypsin, and penicillin-streptomycin were from Invitrogen (Carlsbad, CA). Cell culture dishes were from Sarstedt AG & Co (Nümbrecht, Germany), and 96-well plates were from VWR (Radnor, PA). VTX (unlabelled and tritium labeled) and analogue compounds were kindly provided by H. Lundbeck A/S (Valby, Denmark). [^3^H]-GR65630 was from Perkin Elmer (Waltham, MA). Granisetron and ondansetron were from Tokyo Chemical Industries (Zwijndrecht, Belgium).

### Molecular Biology and Chimera construction

All restriction enzymes were from New England Biolabs (Ipswich, MA). The plasmid vector pcDNA3.1 containing h5HT_3A_ cDNA was kindly provided by B. Niesler (University Hospital Heidelberg Institute of Human Genetics) and was subcloned into the Xenopus oocyte and mammalian cell dual expression vector pXOON as previously reported (Ladefoged, 2018). H. Lundbeck kindly provided pGEM-HE plasmid vectors containing r5-HT_3A_ and gp5-HT_3A_ cDNA. These cDNAs were subcloned into the pXOON vector using EcoR1 and HindIII restriction sites, yielding the plasmid constructs pXOON-r5-HT_3A_ and pXOON-gp5-HT_3A_ (Roche Diagnostics, Mannheim, Germany). Point mutations were generated by site-directed mutagenesis using the QuickChange mutagenesis kit (Stratagene, La Jolla, CA). Successful mutations were verified by DNA sequencing (GATC Biotech, Konstanz, Germany). For the construction of chimeric receptors, a BsrG1 restriction site was introduced by silent mutation at the codons encoding Asp169 and Gln170 in the coding sequence of h5HT_3A_, r5-HT_3A_, and gp5-HT_3A_. Chimeric receptor constructs were generated by excision and swapping of a fragment encoding the region from position 193 to 230 (amino acid numbering refers to the sequence of h5HT_3A_) using the BsrGI site and a native SgrAI restriction site, except for the construction of chimera C7, where a synthetic DNA fragment containing the guinea pig loop C sequence was used (Eurofins Genomics, Ebersberg, Germany). Successful chimera generation was confirmed by sequencing. mRNA was generated using the AmpliCap-Max kit (CellScript, Philadelphia, PA) according to the manufacturer instructions.

### Mammalian Cell Culturing and Expression

Human embryonic kidney 293 (HEK-293) cells (American Type Culture Collection, Manassas, VA) were cultured in growth medium (DMEM supplemented with 10% v/v fetal bovine serum, 100 units/ml penicillin, and 100 μg/ml streptomycin) at 37 °C in a humidified 5% CO_2_ environment. For expression of wild-type (WT) and mutant receptors, HEK-293 cells were transfected using TransIT LT1 DNA transfection reagent (Mirus, Madison, WI) as described previously (Ladefoged, 2018).

FlexStation Assay – FlexStation experiments were performed essentially as described in Price et al. (2005) and Ladefoged et al. (2018). Briefly, the FLIPR blue membrane potential dye (Molecular Devices, Sunnyvale, CA) was diluted in Flex50 buffer (in mM: 57.5 NaCl, 57.5 NMDG+, 1 KCl, 1 MgCl2, 1 CaCl2, 10 HEPES and 10 Glucose; pH 7.4) according to the manufacturer’s instructions. Cells were incubated with 100 µL FLIPR Blue loading buffer per well at 37°C for 30 minutes. Fluorescence was measured in a FlexStation every 2 s for 200 s. At 18 seconds 25 µL of ligand or buffer was added. For VTX concentration inhibition experiments the indicated concentration of VTX was included in the loading buffer during the incubation, and responses were obtained by adding 5-HT (final concentration 30 µM unless otherwise indicated). To account for variation in absolute signal size from different experiments, concentration-response curves were obtained for functional WT and mutant receptors by normalization to the maximum response, or the inhibitor-free response in the case of inhibitor experiments. For the screening of agonist activity of VTX and analogous compounds, a final concentration 1 µM was used unless otherwise indicated. All responses were normalized to the responses to 1 µM 5-HT observed in the same plate.

### Two-electrode voltage clamp electrophysiology

Care and use of live Xenopus laevis were in strict adherence to a protocol (license 2014−15−0201−00031) approved by the Danish Veterinary and Food Administration. Stage V – VI defoliated oocytes were prepared as described previously (Poulsen et al., 2013). Oocytes were injected with either 12 ng of mRNA into the cytoplasm or 0.6 ng of plasmid DNA into the nucleus. Oocytes were stored at 18 C in standard Barth’s solution (in mM: 88 NaCl, 1 KCl, 0.41 CaCl_2_, 2.4 NaHCO_3_, 0.33 Ca(NO_3_)_2_, 0.82 MgSO_4_, 5 Tris (pH 7.4) supplemented with 50 µg/mL gentamicin) and used for two-electrode voltage clamp (TEVC) electrophysiology 3 – 6 or 2 – 5 days after injection of DNA or RNA respectively. Glass pipettes (Glass capillaries, World Precision Instruments) were pulled using a P-1000 micropipette puller (Sutter instruments) such that they had a resistance of 5-30 mΩ. Recording pipettes contained 3 M KCl. Recordings were made with an OC-725C amplifier (Warner Instruments Corp.) and a CED 1401 analogue-digital converter (Cambridge Electronic Design). Oocytes were placed in the recording chamber under continuous superfusion with frogs Ringer’s solution (115 mM NaCl, 2 mM KCl, 5 mM HEPES, 1.8 mM BaCl, pH 7.6) and voltage clamped at −20 mV unless otherwise indicated. All ligands were dissolved in frogs Ringer’s solution and applied using an automated programmable perfusion system (Bioscience Tools). Desensitization rate constants (τ_des_) were determined by fitting to standard single exponential decay functions by least sum of squares using ClampFit 10 software (Molecular Devices). In fast solution switching experiments for testing for current responses, solutions containing saturating concentrations of agonist were applied to the oocyte using a rapid perfusion system consisting of double-barrel glass flow pipes connected to an SF-77 fast solution delivery device (Warner Instruments). This approach allowed rapid switching between control- and agonist-containing solutions. Rapid switching is essential to ensure resolution of receptor currents that desensitize fast. Solution exchange times were determined after each oocyte recording by stepping a dilute external solution across open electrode tips to measure a liquid junction current. The 10–90% rise times for solution exchange were consistently ∼200 ms or less. Programming of perfusion protocols and collection of recordings was performed using WinWCP (Strathclyde University).

### Voltage clamp fluorometry

Voltage clamp fluorometry (VCF) experiments were performed as described in Munro et al. (2018). Briefly, oocytes expressing h5-HT_3A_ Y89C, Q146C or M223C were labelled in 20 µM MTS-TAMRA for 30 seconds in the dark and then stored for up to 3 hours in frog Ringer’s. Recordings were performed using a glass bottom chamber mounted on an inverted fluorescence microscope. The microscope was equipped with a 40x air objective and a G-1B filter set (Chroma 11002v2, excitation filter 541-551 nm, dichroic mirror > 565 nm, emission > 590 nm). Excitation was achieved using a ThorLabs LED (M530L3 Green; nominal wavelength 530 nm). Fluorescence intensity was measured with a D104 photomultiplier tube (PMT) system (Photon Technologies International). PMT voltage was adjusted daily between 450 and 700 volts such that baseline converted fluorescence output on labelled oocytes was between 0.5 and 5 volts.

Radioligand Binding – Radioligand binding experiments were performed essentially as described in Ladefoged et al. (2018) and Thompson et al. (2011). For dissociation experiments, harvested HEK-293 cell membranes expressing h5-HT_3A_ receptors were incubated in 500 µL of 10 mM HEPES buffer (pH 7.4) containing 2 nM [^3^H]-VTX for one hour. HEPES buffer was supplemented with 0.125 % Triton X as this was found to decrease non-specific [^3^H]-VTX binding. Dissociation was initiated by addition of 10 µM unlabelled VTX, and terminated by vacuum filtration onto GF/B filters (Sigma, St. Louis) pre-soaked in 0.3% w/v polyethyleneimine. Radioactivity was determined by scintillation counting using a Beckman BCLS6500 instrument (Fullerton, CA). For competition binding experiments cell membranes expressing h5-HT_3A_ WT or mutant receptors were incubated in 250 µL 10 mM HEPES buffer (pH 7.4) containing 0.3 nM [^3^H]-GR65630 (Perkin Elmer, Waltham, MA) and varying concentrations of VTX or analogues. Following 1 hour of incubation, membranes were captured onto GF/B 96 well plates pre-soaked in 0.3 % w/v polyethyleneimine using a Packard Bell cell harvester. Filter plates were soaked in Microscint-0 scintillation cocktail (Perkin Elmer), and radioactivity was determined using a Packard TopCounter (Packard Inc., Prospect, CT, USA).

### Cultured cells and protein expression for cryo-EM

The mouse wild-type 5-HT_3A_ receptor was expressed in a previously developed stable, inducible HEK-293 cell line from the parental T-Rex 293 cell line (ATCC CRL-1573) (Hassaine et al., 2013; Hassaïne et al., 2017; ^P^olovinkin et al., 2018). The human wild-type 5-HT_3A_ receptor was also expressed in a developed stable, inducible cell line derived from HEK T-Rex 293 cell line (ATCC CRL-1573). The expression construct was cloned in the pcDNA5/TO vector (Gibco) along with sequences encoding four Strep-tag II affinity tags connected by short linkers and followed by a TEV protease cleavage site fused at the N-terminal of the mature protein. The cells were cultured in suspension in FreeStyle 293 medium (Gibco) supplemented with 1% of fetal bovine serum (Gibco), 5 μg/mL blasticidin S (Invivogen) and 100 μg/mL hygromycin B (Invivogen) in an orbital incubator at 37°C with 8% CO_2_.

### Protein production and purification

Cell cultures were expanded until they reached a volume about 5L and a cell density of 2.5×10^6^ cells/mL. The protein expression of the mouse or human 5-HT_3A_ was induced by the addition of 4 μg/mL tetracycline (Sigma). Valproic acid (Sigma) 4 mM final concentration was added 24h post-induction and cells were cultured for another 24-36h in the same orbital incubator at 37°C with 8% CO_2_. Cells were collected by centrifugation (4,000*g* for 20 min at 4°C), the pellets were frozen and stored at −80°C.

The purification of the human 5-HT_3A_ receptor was performed following the same protocol as for the purification of mouse 5-HT_3_ receptor, previously described (Hassaïne et al., 2017; Polovinkin et al., 2018)). Briefly, 30 g of cells were thawed, resuspended in buffer A (10 mM HEPES pH 7.4, 1 mM EDTA, protease inhibitor cocktail, 10 mL of buffer per gram of cells) and mechanically lysed with a T25 digital Ultraturrax disperser (IKA). Membranes were collected by ultracentrifugation (100,000*g* for 1 hour), resuspended in buffer B (50 mM Tris pH 8, 500 mM NaCl, protease inhibitor cocktail, 20 mL of buffer per gram of membrane) and homogenized using a Dounce homogenizer. The solution was supplemented with 0.15% of Anapoe-C12E9 for solubilisation under gentle stirring for 1.5 hours. The insoluble material was removed by ultracentrifugation (100,000*g* for 45 minutes). Solubilized proteins were purified by affinity chromatography using a gravity flow Strep-Tactin resin (IBA, typically 25 mL of resin pre-equilibrated with buffer B supplemented with 0.15% of Anapoe-C12E9). The column was washed with 5 column volumes of buffer C (50 mM Tris pH 7.5, 125 mM NaCl, 0.01% C12E9) and elution was performed using 3 column volumes of buffer C supplemented with 1 mg/mL d-desthiobiotin (Sigma). The elution fraction was concentrated to 0.5 mg/mL using 100-kDa cut-off filters (Millipore) and incubated overnight with homemade enzymes (0.04 mg of TEV protease and 0.1 mg of PNGase F per 1 mg of protein) for affinity tag cleavage and carbohydrate digestion respectively. Thereafter, the protein mixture was concentrated to 1.5 mg/mL and applied on a Superose 6 Increase column (GE healthcare, pre-equilibrated with buffer C for further purification by size-exclusion chromatography. All steps were carried out at 4°C. The most homogeneous fractions of the 5-HT_3A_ receptor were pooled, and concentrated to ∼1 mg/mL for further nanodisc reconstitution or up to 5 mg/mL for grid preparation.

### Nanodisc reconstitution of the human 5-HT3A receptor

The collected concentrated receptors from the previous step (1.5 mg/mL) were mixed with asolectin lipids (Sigma-Aldrich) solubilized at 5 mg/mL in 5% DDM (Anatrace). After 30 minutes incubation, MSP1E3D1(-) (a gift from Stephen Sligar, Addgene plasmid #20066, expressed and purified as previously described (Desinov et al. 2007)) was added to the mixture, which was incubated for 30 additional minutes, before the addition of Bio-Beads (Sigma-Aldrich) at 10 mg/mL final concentration. The molar ratio of the receptor over the MSP and the lipids was 1:7:200. The mixture was incubated under gentle rotation overnight for detergent removal and nanodiscs reconstitution. Bio-Beads were removed by centrifugation (250*g*, 10 min) and the supernatant was subjected to size-exclusion chromatography in a Superose 6 increase column (GE healthcare) equilibrated in SEC buffer (50 mM Tris-HCl, 125 mM NaCl, pH 7.5). All steps were performed at 4°C. The fractions containing the reconstituted receptor in nanodiscs were pooled, concentrated to 1.5 mg/mL, aliquoted, snap frozen in liquid nitrogen and stored at −80°C.

### Electron microscopy and image analysis

Purified mouse or human 5-HT_3A_Rs in detergent or reconstituted in nanodiscs were incubated with 10 µM vortioxetine final concentration for 10 minutes on ice. For the human 5-HT_3A_ receptor in *apo* state no ligand was added. Grids were prepared by depositing 3.5 μl of sample on a glow-discharged (25 mA, 50 s) Quantifoil copper-rhodium or AU/C R 1.2/1.3 grid, blotted at 8°C for 10 s at force 0 using a Mark IV Vitrobot (Thermo Fisher) and plunge-frozen in liquid ethane. The datasets were recorded on a Titan Krios Cryo-TEM (Thermo Scientific) at the European Synchrotron Radiation Facility (ESRF) (Kandiah et al., 2019) or on a Glacios Cryo-TEM (Thermo Scientific) at the Institut de Biologie Structurale (IBS), as detailed in Table SI3.

Picking was performed with Cryolo. The rest of the data processing was performed in Relion or Cryosparc using standard procedures. Typically, the bad micrographs were excluded using manual inspection with Curate exposure; rounds of 2D classification permitted to select class averages that resembled a pentameric ion channel. Rounds of Ab-initio model generation followed by heterogenous refinements were used to select the final sets of particles. The 3D reconstructions were then obtained after a round of non-uniform refinement (followed by local refinement in the case of C1-refined sets). Details about the workflow for each dataset are presented in Fig. SI3, SI7, SI8 and SI9.

### Computational modeling and ligand docking

For the mouse receptor, the cryo-EM structure in complex with VTX was used as an initial template for molecular-dynamics simulations. For the human receptor, a composite model encompassing the cryoEM structure in complex with VTX, for the ECD, and the apo active/distorted cryoEM structure, for the TMD, was used. The models were inserted in a fully hydrated phospholipid bilayer using CHARMM-GUI (Jo et al., 2008; Wu et al., 2014). The human and the mouse molecular assays resume to initial boxes of ∼130 x 130 x 143 Å^3^ and ∼130 x 130 x 190 Å^3^ of ∼222,000 and ∼284,0000 atoms, composed of 350 POPC (1-palmitoyl-2-oleoyl-sn-glycero-3-phospho-choline), and ∼48,000 and ∼68,000 water molecules, respectively. Both systems were neutralized and brought to a salt concentration of 0,150 M NaCl.

Proteins, lipids and ions were modeled using the CHARMM36 force field (Best et al., 2012; Klauda et al., 2010). Water was described by the TIP3P model (Jorgensen et al., 1983). VTX was modeled using the CHARMM-like force-field parameters derived previously (Ladefoged et al., 2018).

All molecular-dynamics (MD) simulations were performed with NAMD3 (Phillips et al., 2020) in the isobaric-isothermal ensemble at a pressure of 1 atm and a temperature of 300 K. The temperature was controlled using Langevin dynamics with a damping coefficient of 1 ps^-1^. The pressure was kept constant using a semi-isotropic modified Nosé-Hoover algorithm in which Langevin dynamics is used to control the barostat fluctuations (Feller et al., 1995). Long-range interactions were treated by means of the particle-mesh-Ewald procedure (Darden et al., 1993). All bonds involving hydrogen atoms were constrained to their nominal force-field length using the SETTLE algorithm (Miyamoto and Kollman, 1992) for water molecules and RATTLE (Andersen, 1983) for all other bonds. A hydrogen-mass-repartitioning scheme (Balusek et al., 2019), wherein hydrogen masses were scaled up threefold and the masses of adjacent atoms were reduced accordingly, was employed allowing integration of the equation of motion with a 4fs time step. The RESPA multiple-time-step integration scheme (Tuckerman et al., 1992) was also used where the short-range interactions were updated at each time step and the slow varying long-range interactions every two time steps.

MD trajectories for both systems were produced using the following protocol. First the energy of the full system was minimized during 200 steps with all non-hydrogen-protein atoms being harmonically restrained to their initial positions. Next, the system was thermalized during 50 ns with all-backbone atoms restrained to their initial positions. The harmonic restraints were removed step-wisely over a period of 4 ns. Starting from the equilibrated configuration, five independent trajectories of 1 µs were generated and subsequently employed for statistical analysis. Pore-profile averages were determined using VMD (Humphrey et al., 1996) and the HOLE program (Smart et al., 1996).

### Data and Statistical Analysis

The data and statistical analysis comply with the recommendations on experimental design and analysis in pharmacology (Curtis et al., 2015). All data analysis was performed using GraphPad Prism. Data is given as either mean ± SD or mean with 95% confidence interval [95% CI] unless otherwise indicated. IC_50_ and EC_50_ curves were fit to the equation

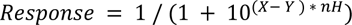

where *Response* is the agonist-evoked response measured at a given inhibitor concentration normalized to the response in either absence of inhibitor (for IC_50_) or the maximum response (for EC_50_), *Y* is the concentration of inhibitor (for IC_50_) or agonist (for EC_50_) that produces a half-maximal inhibition or response, *X* is the logarithm of the concentration of the inhibitor or agonist concentration, and *n_H_* is the Hill slope. Competition binding experiments were fit to the Cheng-Prusoff equation (Cheng and Prusoff, 1973) used based on the assumption that VTX and the radioligand bind competitively under steady-state conditions.

### Resource availability

Further information and requests for resources and reagents should be directed to and will be fulfilled by the Lead Contact, Dr. Anders S. Kristensen.

### Materials availability

Plasmids generated in this study are available from the lead contact upon request.

---

**Fig. SI1.**
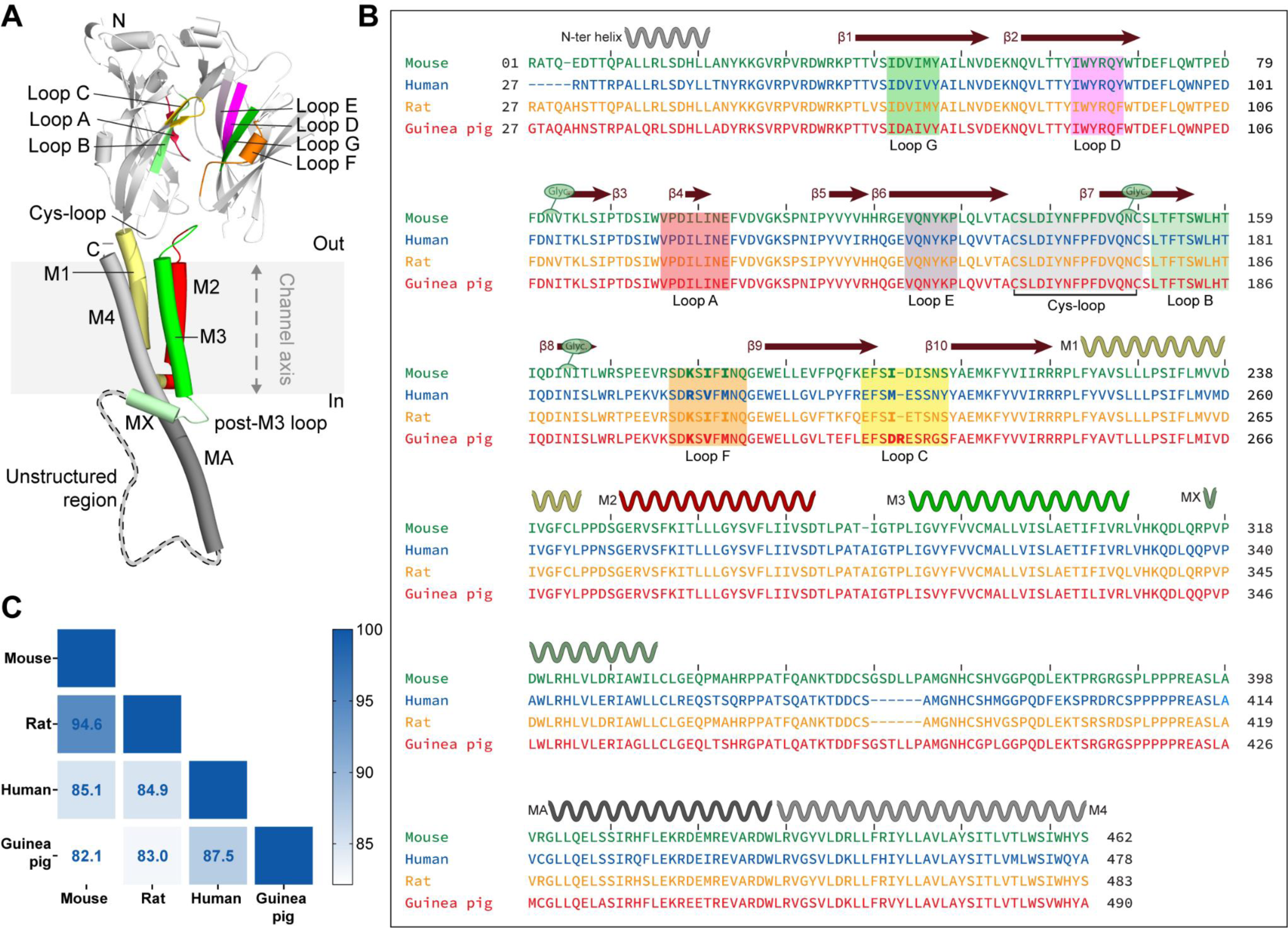
Nomenclature, sequence alignment. **A.** Ribbon illustration of a full 5-HT3R subunit and the extracellular domain of a complementary subunit. Binding elements are indicated in colors: Loops A-C on the principal subunit, three strands and a loop D-G on the complementary subunit. **B.** Multiple sequence alignment of the mouse, human rat and guinea pig 5-HT3A receptors. Binding elements and Cys-loop receptors are highlighted. Residues of interest in Loops F and C are shown in bold. **C.** Amino acid percent identity matrix of mouse, human, rat and guinea 5-HT3A receptors.

**Fig. SI2.**
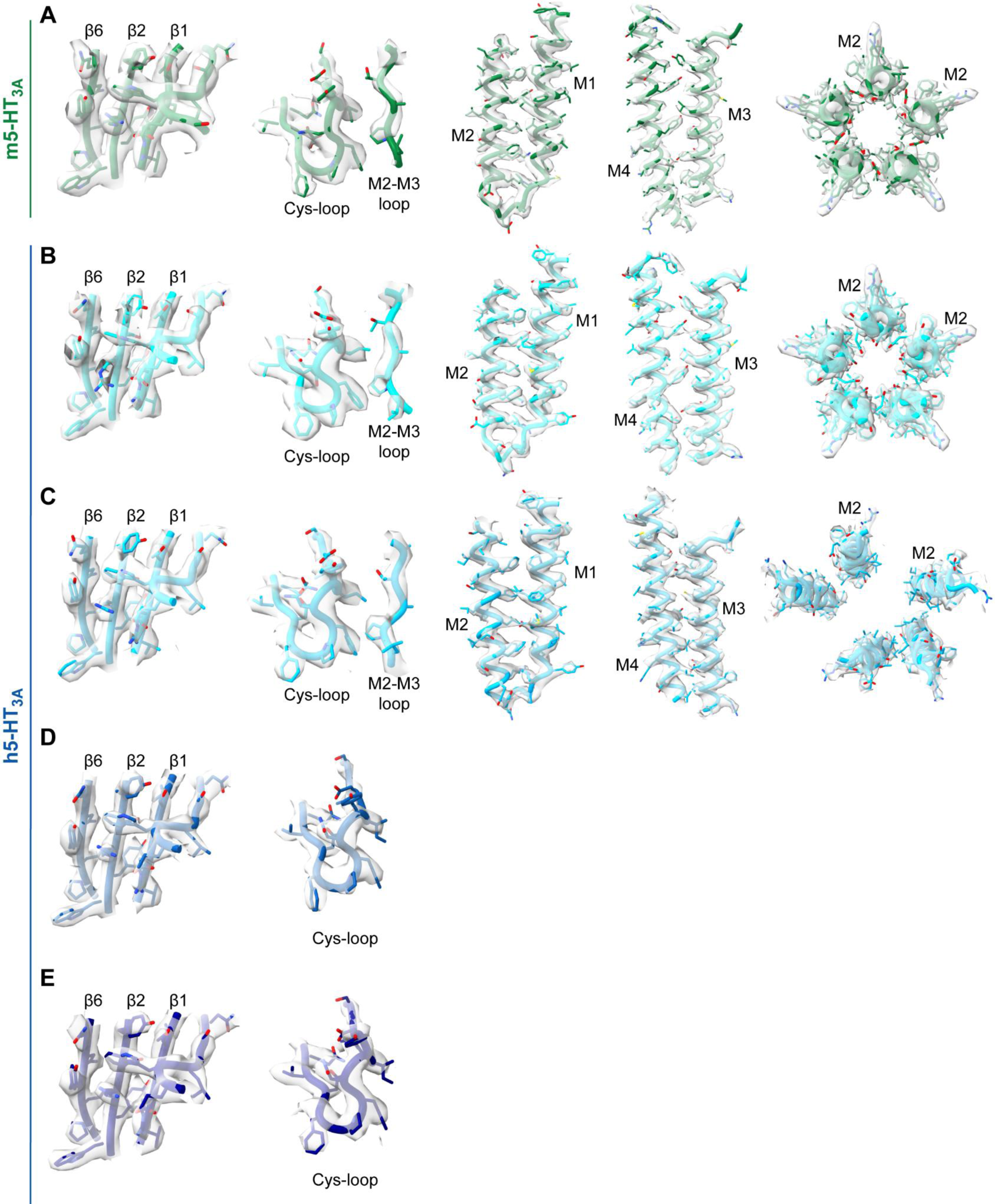
Quality of the density maps. Densities of the **A.** m5-HT3AR (green) and h5-HT3AR (shades of blue) **B.** APO resting, **C.** APO Active-distorted, **D.** in complex with VTX (detergent) and **E.** in complex with VTX (nanodisc) reconstructions in surface representation overlaid with the structure. From left to right: densities of the β-sheets in the ECD, densities of the Cys-loop and/or the M2–M3 loop, densities of helices M1 and M2, densities of M3 and M4, densities of M2 at the level of L9′ (L260).

**Fig. SI3.**
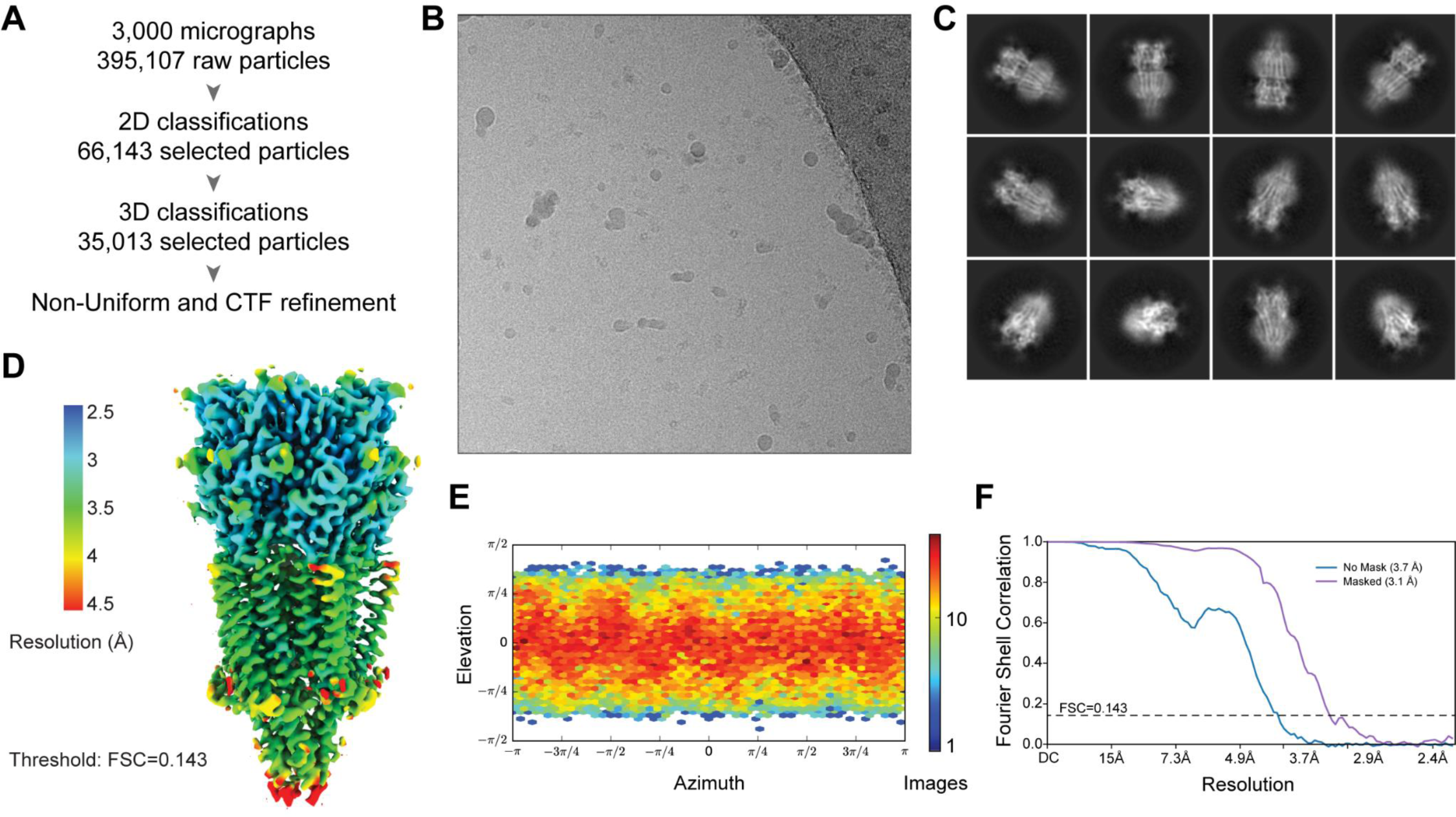
Image analysis workflow, m5-HT3AR in detergent dataset. **A.** Schematic of the image analysis workflow **B.** A representative micrograph of the dataset. **C.** Selected 2D class averages of the final particles set. **D.** Side view of the final reconstruction. The sharpened 3D density map is colored according to the local resolution (FSC threshold of 0.143). **E.** Heat map of the angular distribution of particle projections for the reconstruction. **F.** Gold-standard FSC curves. The dotted line represents the 0.143 FSC threshold.

**Fig. SI4.**
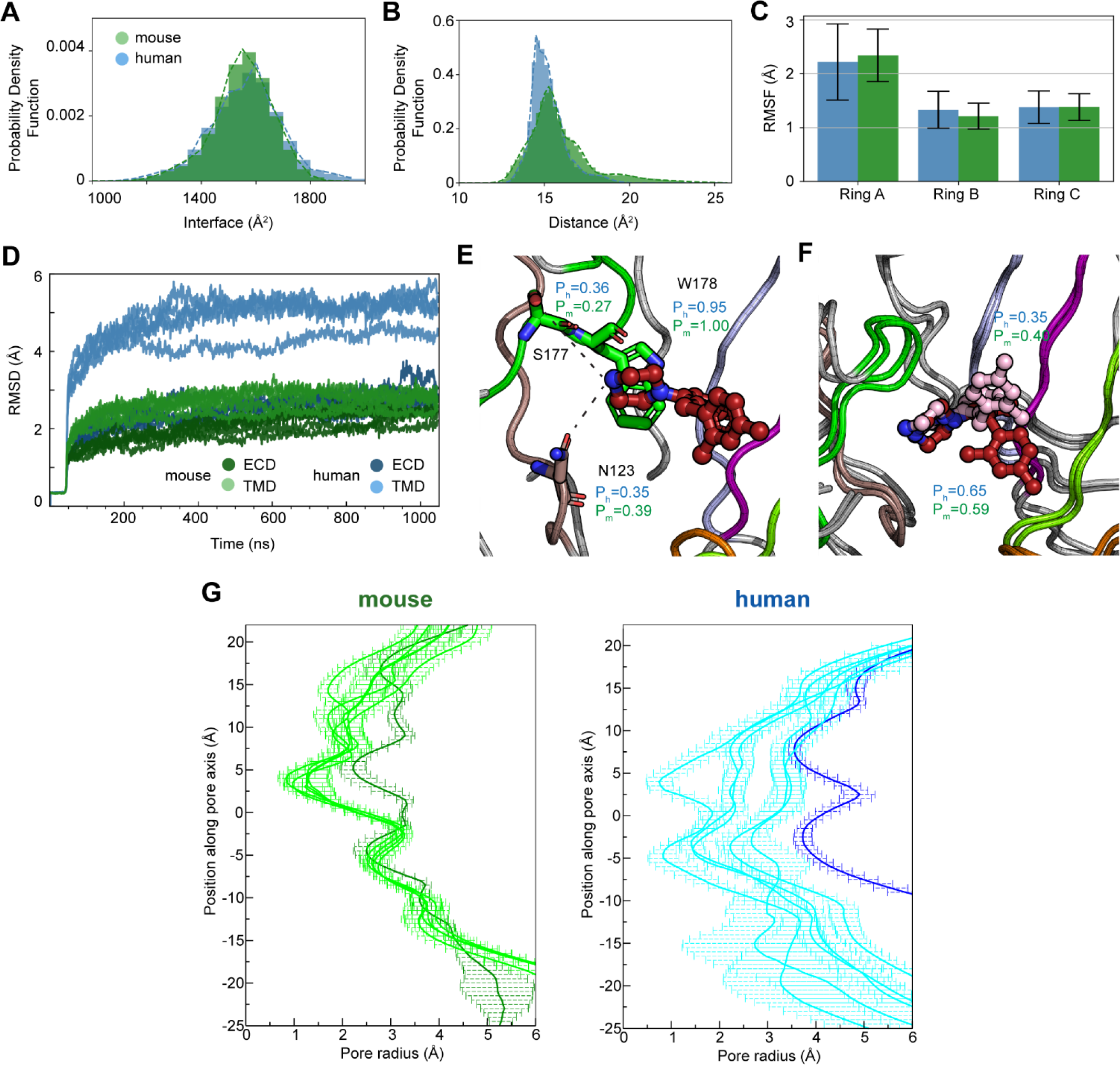
Compartive features of the m5-HT3AR and h5-HT3AR during molecular dynamics simulations. **A.** Probability density function of the surface area buried in ECD-ECD interfaces, for the human receptor (blue) and the mouse receptor (green). The represented data takes into account all 5 interfaces for 5 independent runs. **B.** Probability density function of loop C to loop D distance, for the human receptor (blue) and the mouse receptor (green). The computed distance is between the center-of-mass of loop C Ca atoms and the Cα of the conserved arginine of loop D (residues 222-228 and R87 in h5-HT3AR). In panels A and B, probability density functions (dashed lines) were obtained by fitting a Gaussian mixture model to the distribution pooled across simulation repeats. The number of Gaussian components was determined by the Bayesian information criterion.**C.** Root mean square fluctuations of each of the three vortioxetine cycles, for the human receptor (blue) and the mouse receptor (green). **D**. Time course of the root mean square deviations of Cα atoms, for the human receptor (blue) and the mouse receptor (green). The pale color curves correspond to the TMDs while the dark color curves correspond to the ECDs. The larger deviations for the human receptor TMD is clearly apparent. **E.** Probability of hydrogen bond or cation/π formation between VXT’s primary amine and nearby residues N123, S177, and W178 (human residue numbering) calculated based on the simulations in either human or mouse 5-HT3AR (Ph and Pm, respectively). The probability is calculated based on the calculated distance distribution pooled across simulation repeats (shown data is based on the 500-1000 ns interval of the trajectories, but calculations for 800-900 ns and 900-1000 ns are near-identical). **F**. Probability of phenyl A being in an upwards- or downwards facing conformation in m and h5-HT3AR. The probabilities are calculated the same way as described for E. **G.** Pore profiles during MD simulations. The average over the last 50 ns and the standard deviation are shown for 5 independent runs.

**Fig. SI5.**
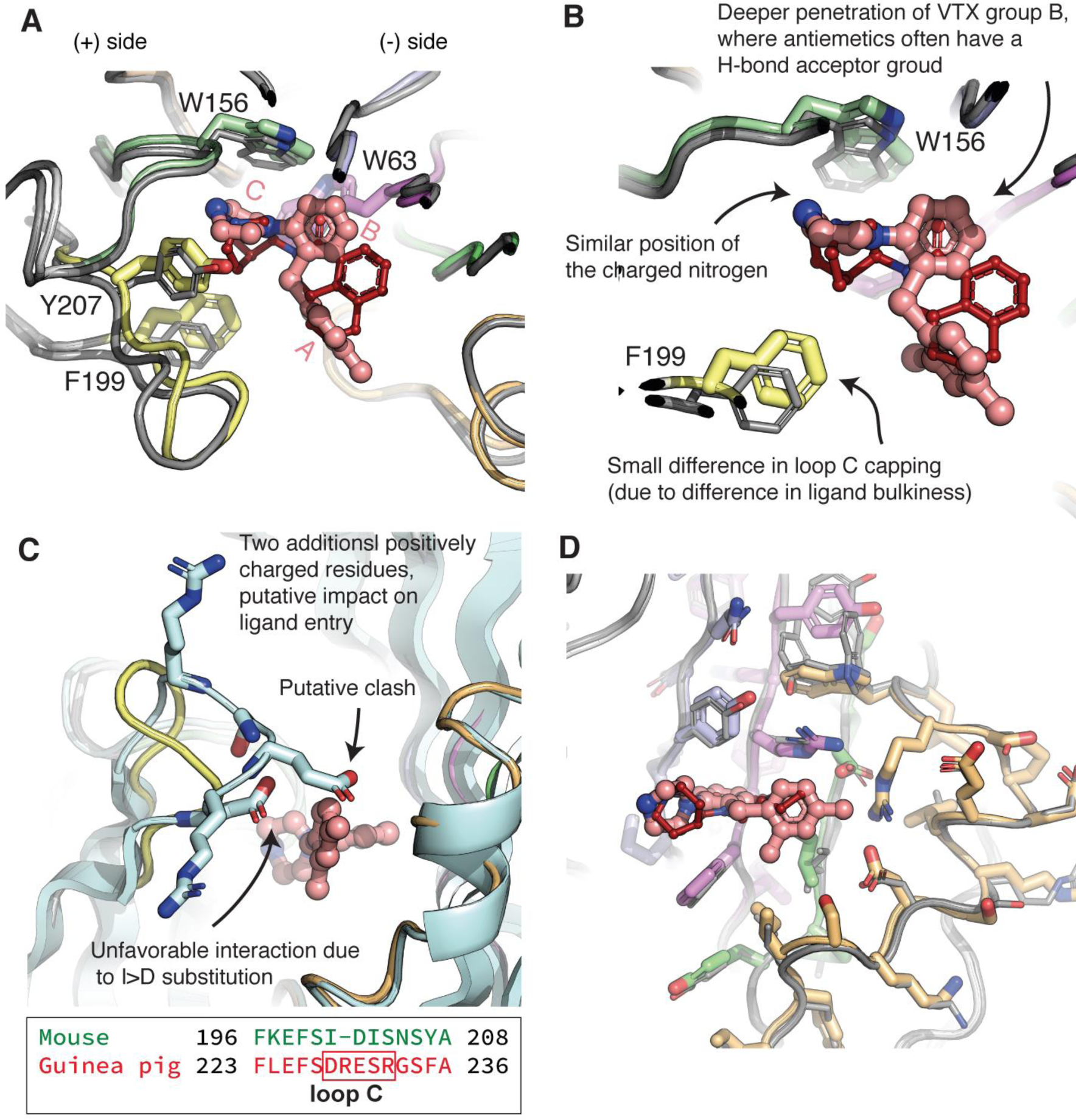
Overlay of the mouse VTX-bound and palonosetron-bound structures. **A.** Overlay of the m5-HT3AR structures bound to VTX (colored as in Fig. 2) and to palonosetron (6Y1Z, gray). Complementary subunits ECDs were used for superimposition. A subset of the aromatic cage residues and the ligands are shown as sticks. The view is similar to that of Fig. 2E. **B.** Zoom on the overlaid ligands, showing the similar position of the charged nitrogen atom interacting with W156, the deeper penetration of the central phenyl group of VTX compared to palonosetron. A difference of position of F199 in loop C is also seen. **C.** Overlay of the m5-HT3AR structures bound to VTX and of the Alphafold model for the guinea pig receptor. The residues of the tip of loop C of the guinea pig model are shown as sticks. Note the unfavorable interaction of D227 with VTX, the putative clash of E229 with the complementary subunit, the presence of two additional charged residues R228 and R231. **D.** Overlay of the m5-HT3AR structures bound to VTX (colored as in Fig. 2) and to palonosetron (6Y1Z, gray), complementary subunit. The resemblance of the VTX-bound structure with those of -setron-bound receptors extends to the organization of side chains neighboring the ligand, on the complementary subunit side. The fact that this half of the binding site has a conserved, rigid structure, structured by strong interactions seems constitutive of the 5-HT3A receptor. Indeed, that organisation is also seen in serotonin-bound structures representative of an active state. Diverse chemical structures thus fit in with only little accommodation of the binding site moiety located on the complementary subunit. The largest differences between structures harboring different ligands lies in deviations of loop C, and more importantly in subunit-subunit interface reorganization (Polovinkin et al., 2018; Zarkadas et al., 2020). In other words, the quaternary structure re-arrangement dominates over tertiary structure deviations. Still the position of the backbone of loop C shows the biggest deviation among the VTX- and setron-bound structures, in line with the classical notion that it acts as a flexible lid on the orthosteric site of every pLGIC.

**Fig. SI6.**
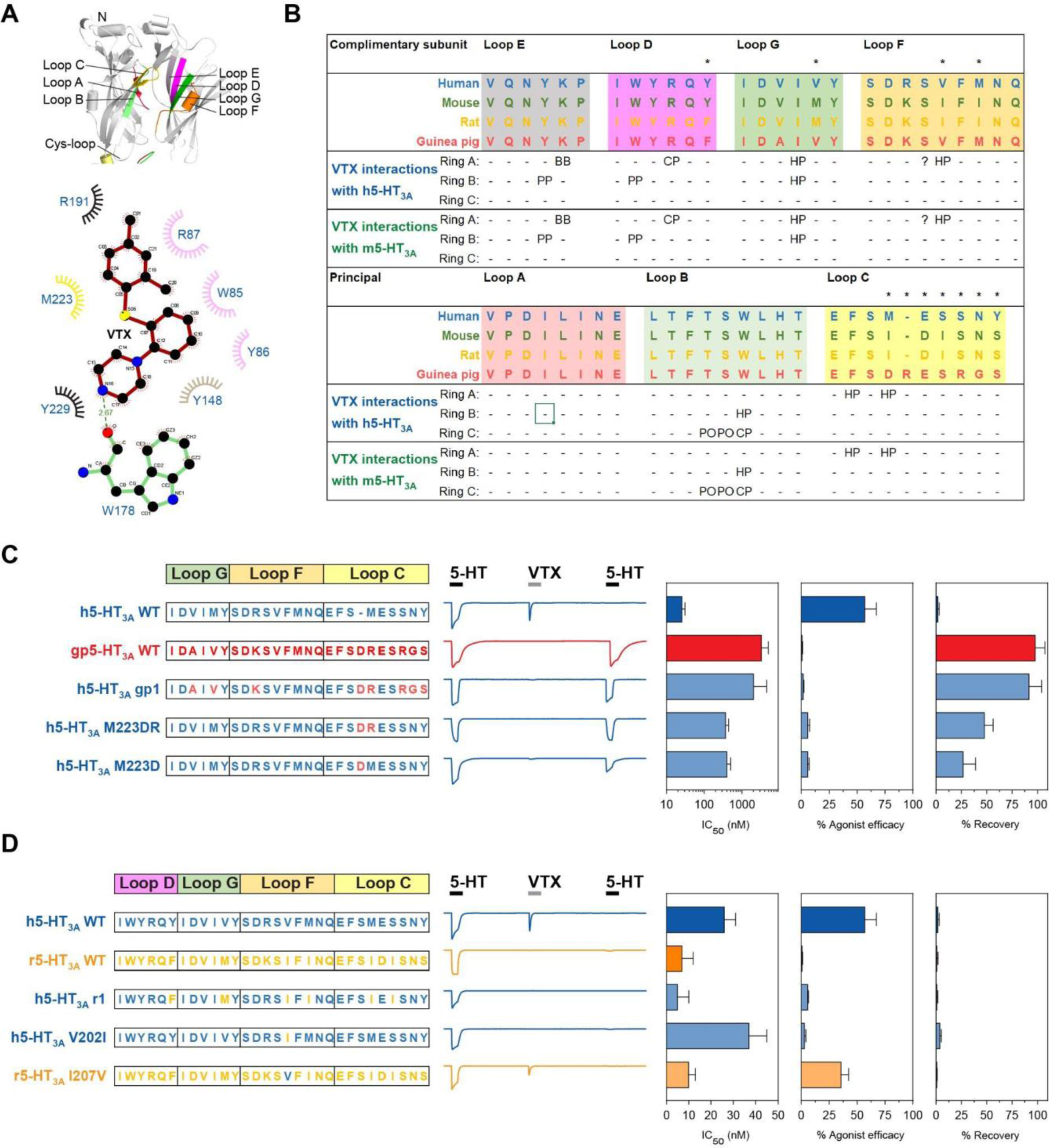
Comparison of VTX binding site residues and ligand-interactions in human and rodent receptors and mutational analysis of determinants for VTX activity. **A.** Overview of the loop segments that form the orthosteric binding site in the 5-HT3A receptor (upper panel) and the 2D ligand-protein interaction diagram of vortioxetine (lower panel). Binding elements are indicated in colors: Loops A-C on the principal subunit, three strands and a loop D-G on the complementary subunit. **B.** Multiple sequence alignment of the loop sequences in mouse (green), human (blue), rat (orange), and guinea pig (red) 5-HT3A receptors. Non-conserved positions are indicated with *. Presence and type of direct molecular interaction with VTX ring A, B, and C are indicated below the alignment: Hydrophobic = HP, hydrogen bond = HB, cation/pi=CP; pi/pi=PP, polar=PO. **C-D.** Amino acid sequence of loops G, F, and C in wt and mutant human, guinea pig and rat 5-HT3A receptors with corresponding representative traces of the current-response to the 3-step sequential protocol applying twice 10 μM 5-HT and 10 μM VTX in between that defines the receptor response-phenotype. To the right is shown a summary of the VTX phenotype for the constructs. Data represent the mean.

**Fig. SI7.**
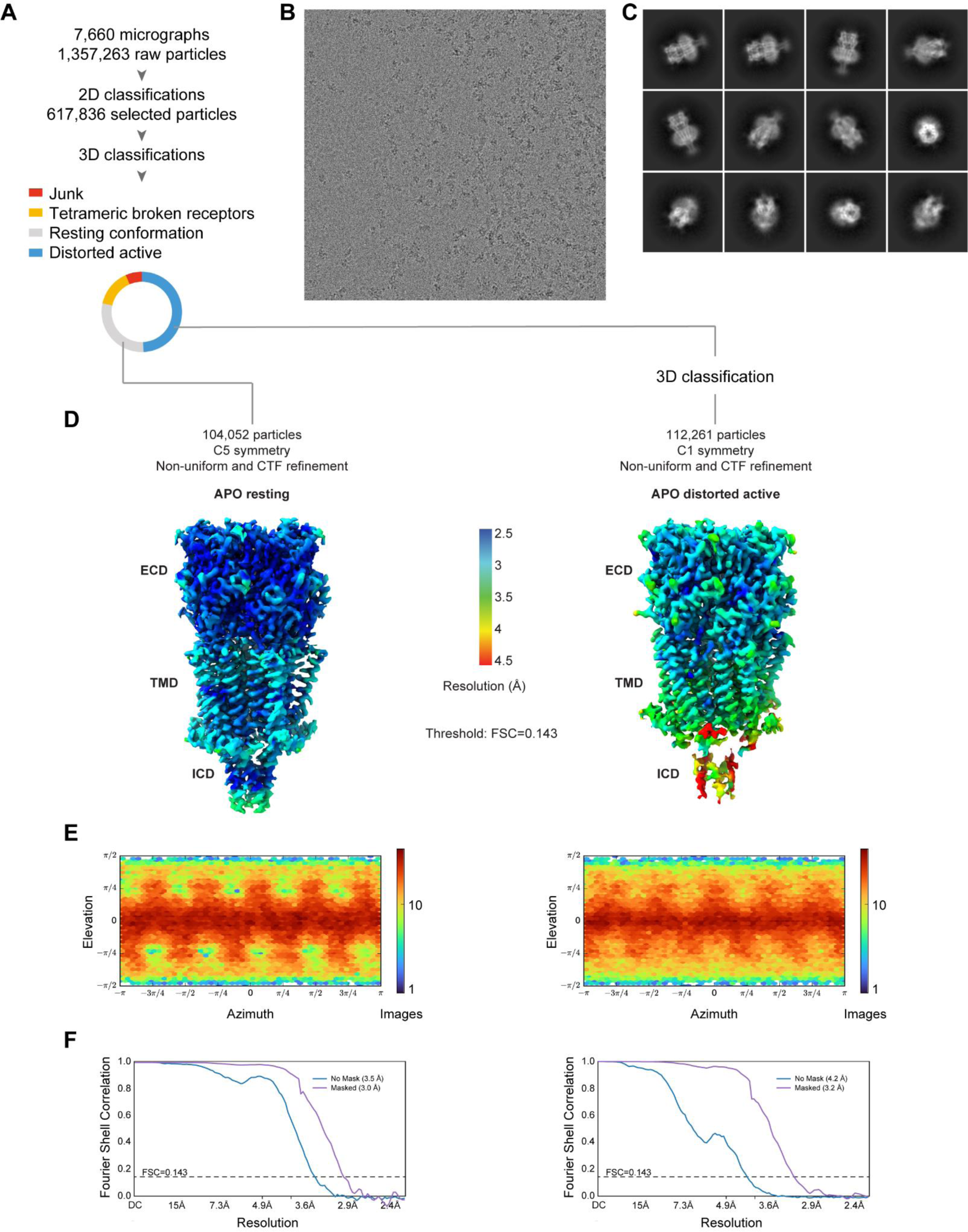
Image analysis workflow, h5-HT3AR in detergent APO dataset. **A.** Schematic of the image analysis workflow **B.** A representative micrograph of the dataset. **C.** Selected 2D class averages corresponding to pentameric receptors. **D.** Side views of the reconstructions used for model building of the APO resting (left) and the APO distorted active conformations (right). The density-modified map is colored according to the local resolution (FSC threshold of 0.5). **E.** Heat map of the angular distribution of particle projections for the reconstructions. **F.** Gold-standard FSC curves. The dotted line represents the 0.143 FSC threshold.

**Fig. SI8.**
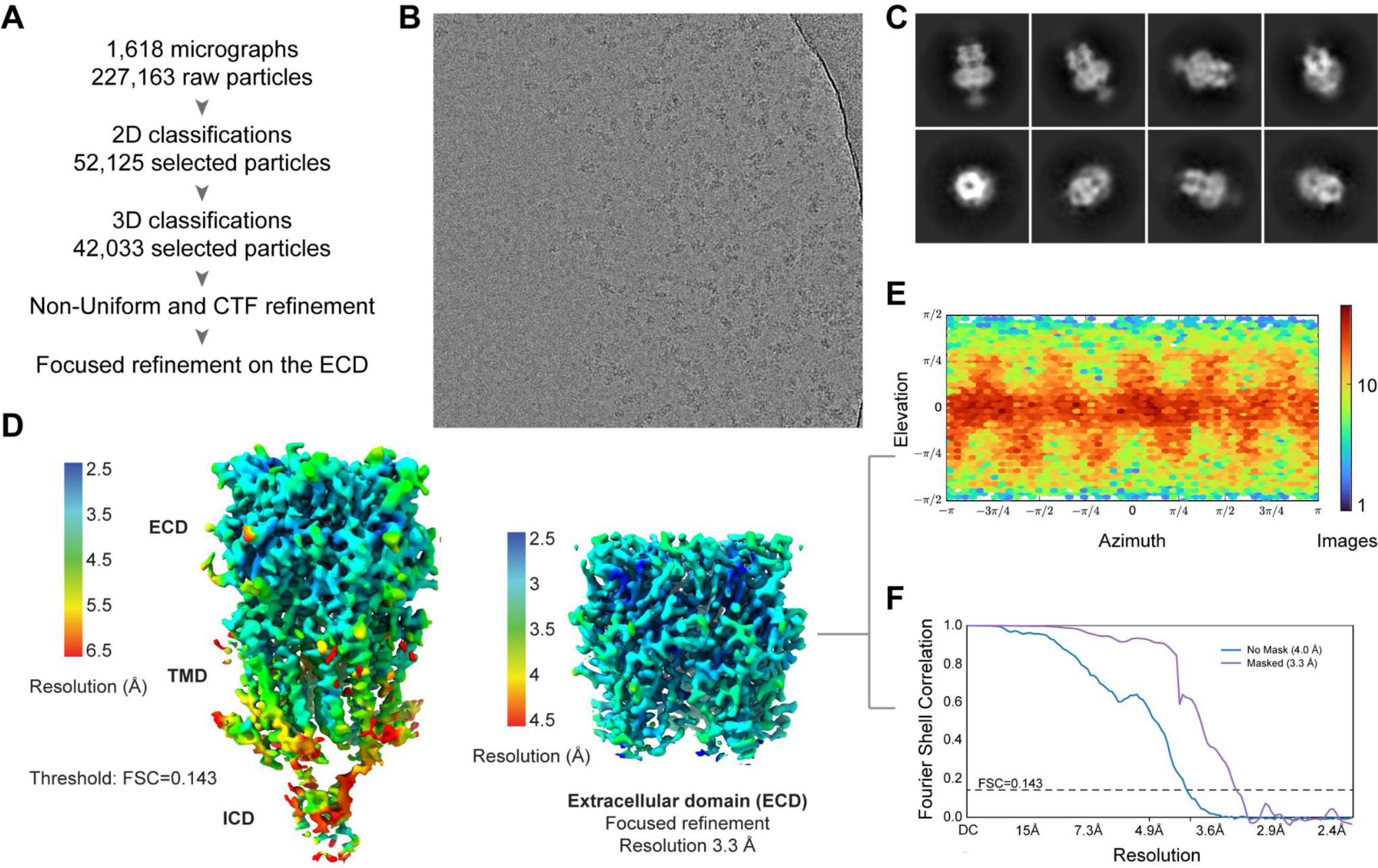
Image analysis workflow, h5-HT3AR in detergent in complex with VTX dataset. **A.** Schematic of the image analysis workflow **B.** A representative micrograph of the dataset. **C.** Selected 2D class averages of the final particles set. **D.** Side views of the final reconstructions for the whole map (left) and for the focused one on the extracellular domain (right). The sharpened 3D density maps are colored according to the local resolution (FSC threshold of 0.143). **E.** Heat map of the angular distribution of particle projections and, **F.** Gold-standard FSC curves for the extracellular domain reconstruction. The dotted line represents the 0.143 FSC threshold.

**Fig. SI9.**
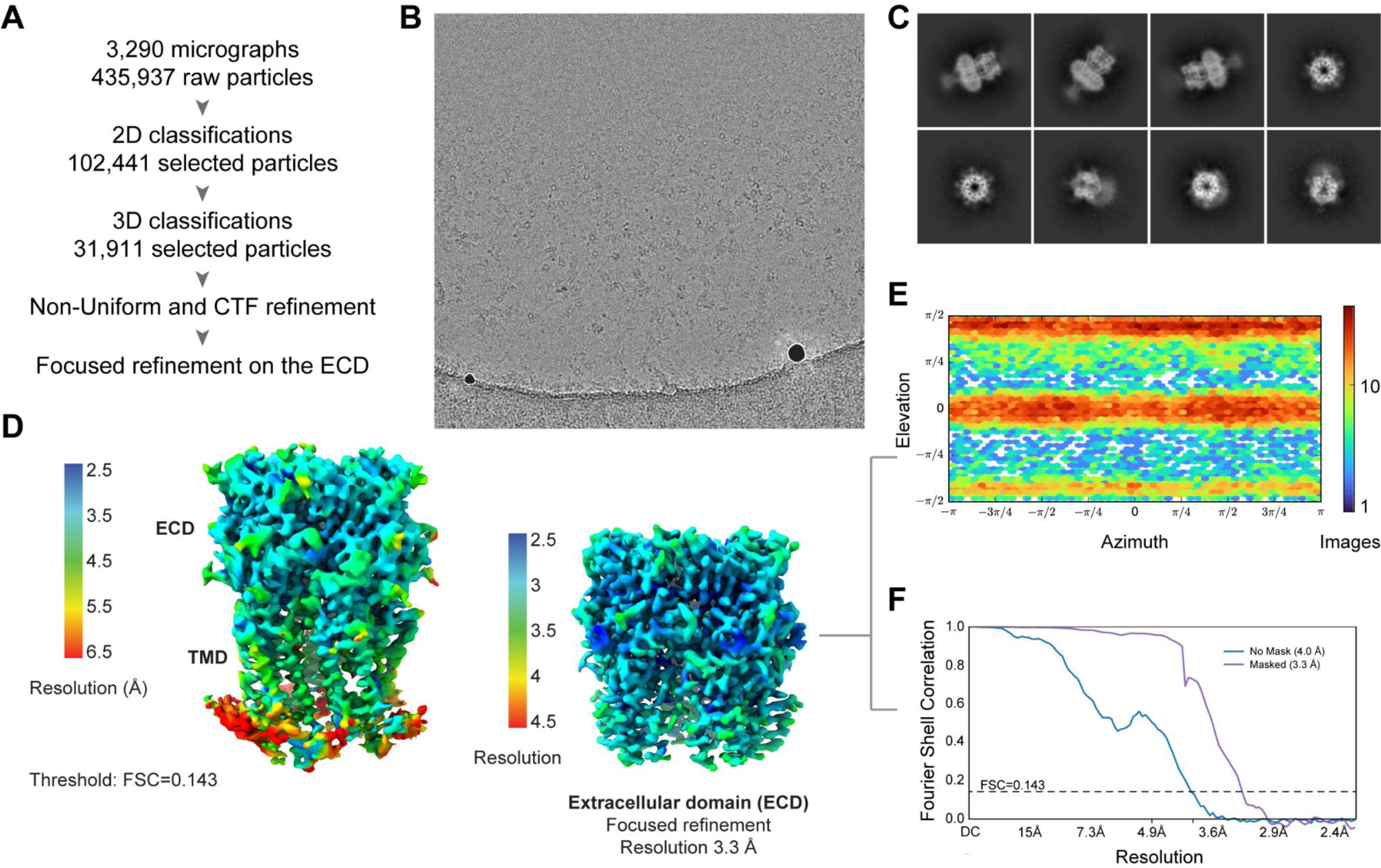
Image analysis workflow, nanodisc reconstituted h5-HT3AR in complex with VTX dataset. **A.** Schematic of the image analysis workflow **B.** A representative micrograph of the dataset. **C.** Selected 2D class averages of the final particles set. **D.** Side views of the reconstruction used for model building. The density-modified map is colored according to the local resolution (FSC threshold of 0.5). One view shows the map around the built model, while the second view shows the whole map at the same contour, highlighting spurious density at the level of the detergent belt and in the ICD. As discussed in the text, the TMD is flexible and probably in a non-physiological conformation. **E.** Heat map of the angular distribution of particle projections for the reconstruction. **F.** Gold-standard FSC curves. The dotted line represents the 0.143 FSC threshold.

**Fig. SI10.**
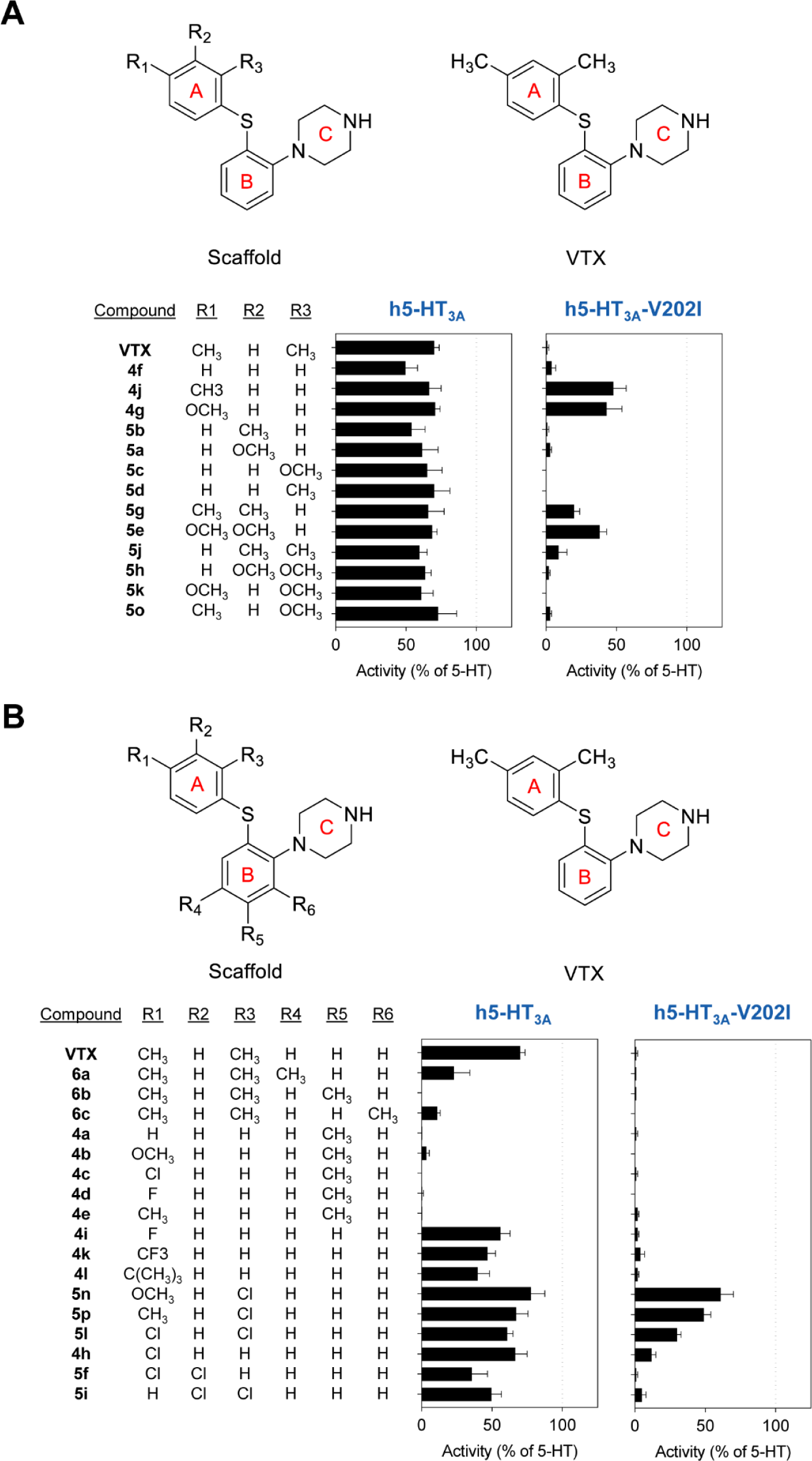
Structure-activity relationship of the agonist effects of VTX analogue compounds. Analogues were tested for agonist response at a concentration of 1 µM in the Flexstation membrane potential assay at the wild-type h5-HT3A and rat-like h5-HT3A-V202I mutant receptors. (A-B) Chemical structures of VTX and analogs with substitutions only in the R1-R3 positions (A) or in the R1-R6 positions (B) along with graphical summary of agonist activity. Data represent the mean ± SEM of the fluorescence response (normalized to the 5-HT response measured in parallel) in least five wells.

**Fig. SI11.**
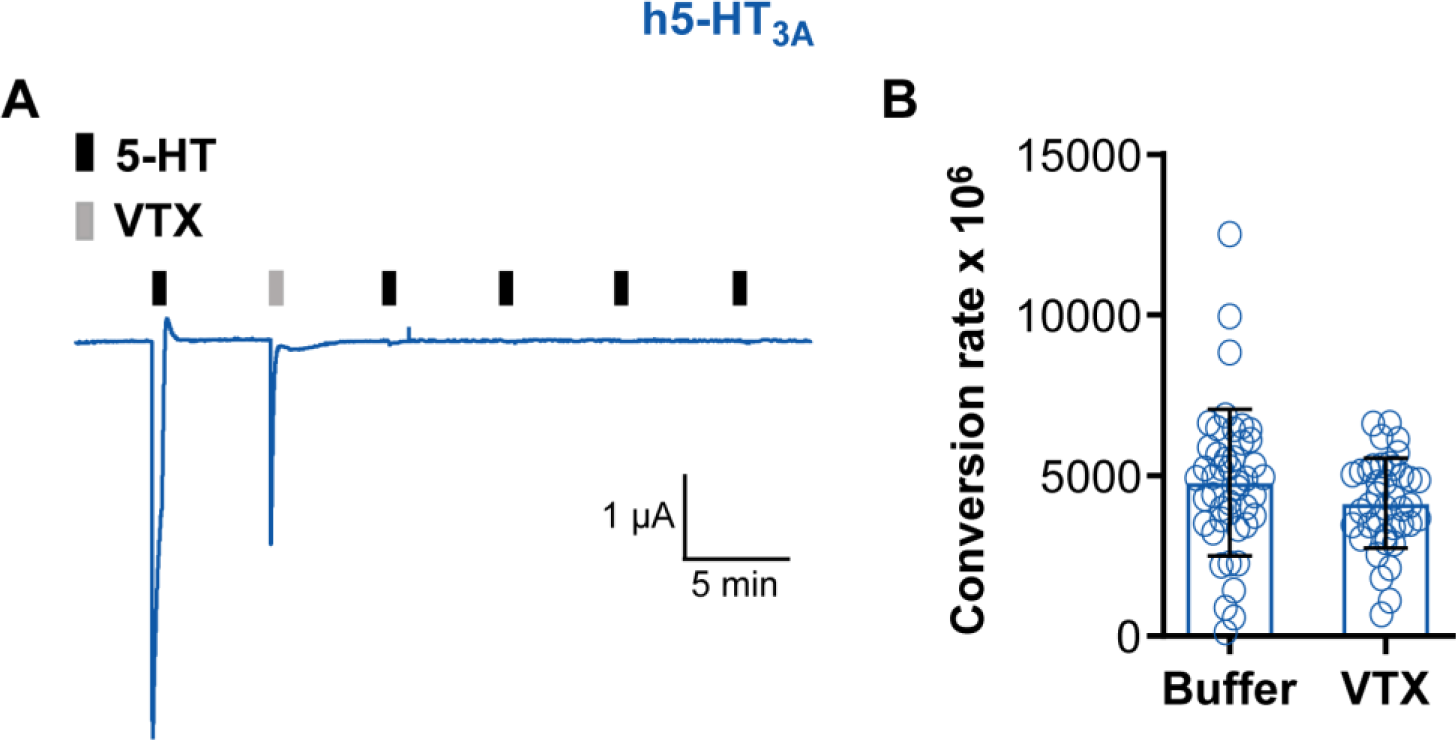
β-lactamase assay. **A.** Representative trace indicating that the persistent inhibition by VTX is maintained in 5-HT3A- β-lactamase construct expressed in Xenopus oocytes. Black bars indicate application of 10 µM 5-HT and the grey bar indicates application of 10 µM VTX. **B.** Representative single day experiment showing lactamase conversion rate of nitrocefin for oocytes expressing 5-HT3A-β-lactamase incubated with either buffer or VTX for 30 minutes.

**Fig. SI12.**
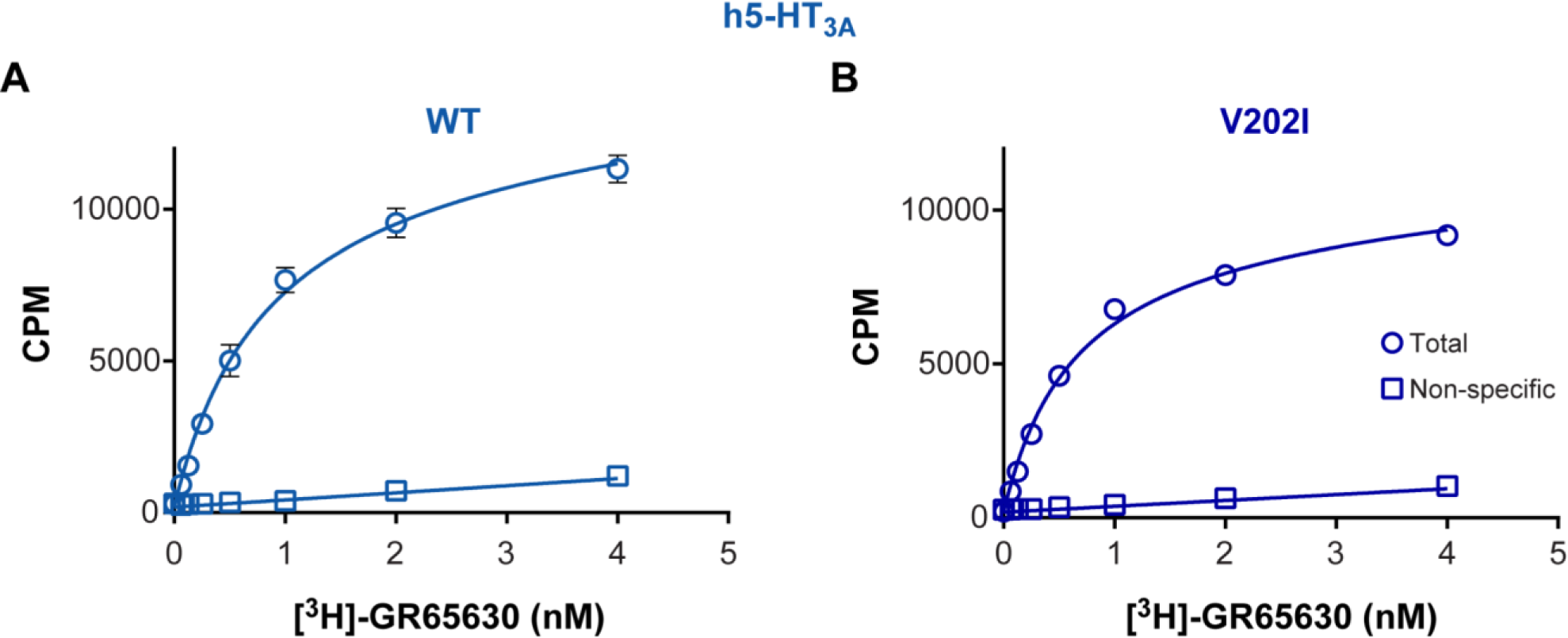
Saturation binding curves for [3H]-GR65630 at WT h5-HT3AR and h5-HT3AR-V202I. **A-B.** Specific binding (circles) and non-specific (squares) binding of [^3^H]GR65630 to membranes from HEK293 cells expressing WT (A) and V202I mutant (B) human 5-HT3AR is shown. The incubation time was 1 h at room temperature in 10 mM HEPES buffer, pH 7.4. Granisetron (1 µM) was used to define non-specific binding. Data points represent the mean + SEM of 3 determinations. The full line is the fit of equation [RL] = [L] Bmax/([L] + *K*d) where [RL] is the specific binding, [L] is the concentration of [^3^H]GR65630, Bmax is the maximum binding capacity, and *K*d is the dissociation constant. The estimated *K*d was 0.82 ± 0.06 nM and 0.62 ± 0.05 nM for WT and V202I, respectively.

**Table SI1.** EC50 and IC50 values. 5-HT EC50 values and VTX IC50 values (determined by FlexStation membrane potential assay using 30 µM 5-HT); and agonism induced by 10 µM VTX relative to 10 µM 5-HT, as well as recovery of responses following 6 minutes of washing after VTX application (determined by Xenopus oocyte electrophysiology) for WT and mutant receptor constructs.

| Receptor | Mutant | 5-HT EC<sub>50</sub> [95% CI] nM | N | VTX IC<sub>50</sub> [95 % CI] nM | N | Agonism (%) | Recovery (%) | N |
| --- | --- | --- | --- | --- | --- | --- | --- | --- |
| h5-HT<sub>3A</sub>R | WT | 193 [175;213] | 28 | 20 [18;23] | 6 | 57 ± 27 | 1 ± 1 | 7 |
| | gp1 | 1091 [948;1256] | 6 | 2061 [955;4445] | 3 | 2 ± 1 | 92 ± 32 | 7 |
| | gp2 | 252 [218;291] | 6 | 403 [324;501] | 7 | 6 ± 3 | 27 ± 32 | 7 |
| | M223D | 306 [208;450] | 6 | 375 [320;438] | 7 | 6 ± 4 | 48 ± 22 | 7 |
| | M223DR | 1049 [859;1280] | 3 | 1011 [659;1550] | 3 | 0 ± 0 | 58 ± 25 | 7 |
| | r1 | 43 [31;58] | 3 | 5 [2;10] | 3 | 6 ± 2 | 1 ± 2 | 7 |
| | R200K | 670 [568;789] | 6 | 94 [81;110] | 5 | 37 ± 6 | 2 ± 2 | 7 |
| | V202I | 251 [208;303] † | 4 | 37 [30;45] † | 3 | 3 ± 3 | 4 ± 3 | 7 |
| | M204I | 535 [397;720] | 5 | 73 [60;89] | 5 | 43 ± 10 | 2 ± 3 | 7 |
| | M223I | 120 [100;130] † | 3 | 20 [16;25] † | 5 | 46 ± 11 | 8 ± 14 | 7 |
| | S225T | 716 [614;834] | 5 | 58 [47;70] | 5 | 44 ± 16 | 4 ± 4 | 7 |
| | Y228S | 112 [80;156] | 4 | 84 [66;106] | 4 | 51 ± 8 | 1 ± 1 | 7 |
| gp5-HT<sub>3A</sub>R | WT | 255 [237;275] | 14 | 2348 [1960;2814] | 6 | 1 ± 1 | 144 ± 41 | 7 |
| r5-HT<sub>3A</sub>R | WT | 1665 [1081;2563] | 15 | 26 [19; 36] | 6 | 1 ± 1 | 2 ± 1 | 7 |
| | I207V | 778 [504;1202] | 11 | 10 [8;13] | 5 | 36 ± 17 | 1 ± 1 | 7 |

**Table SI2.** IC50 values for VTX, ondansetron, and granisetron. IC50 values were determined, using 30 µM 5-HT, at human, rat, and guinea pig 5-HT3A receptors by FlexStation membrane potential assay. The data for ondansetron and granisetron at the human receptor is reproduced from (Ladefoged et al., 2018).

| Species | Inhibitor | IC<sub>50</sub> [95 % CI] nM | N |
| --- | --- | --- | --- |
| Human | VTX | 20 [18 ;23 ] | 28 |
|  | Ondansetron | 7.6 [5.9 ; 9.7] | 7 |
|  | Granisetron | 0.78 [0.58;1.05] | 4 |
| Guinea Pig | VTX | 2348 [1960;2814] | 12 |
|  | Ondansetron | 9810 [7209;1335] | 5 |
|  | Granisetron | 1257 [1069;1477] | 5 |
| Rat | VTX | 26 [19;36] | 10 |
|  | Ondansetron | 18 [2.0;27] | 6 |
|  | Granisetron | 0.15 [0.11;0.20] | 4 |

**Table SI3.**
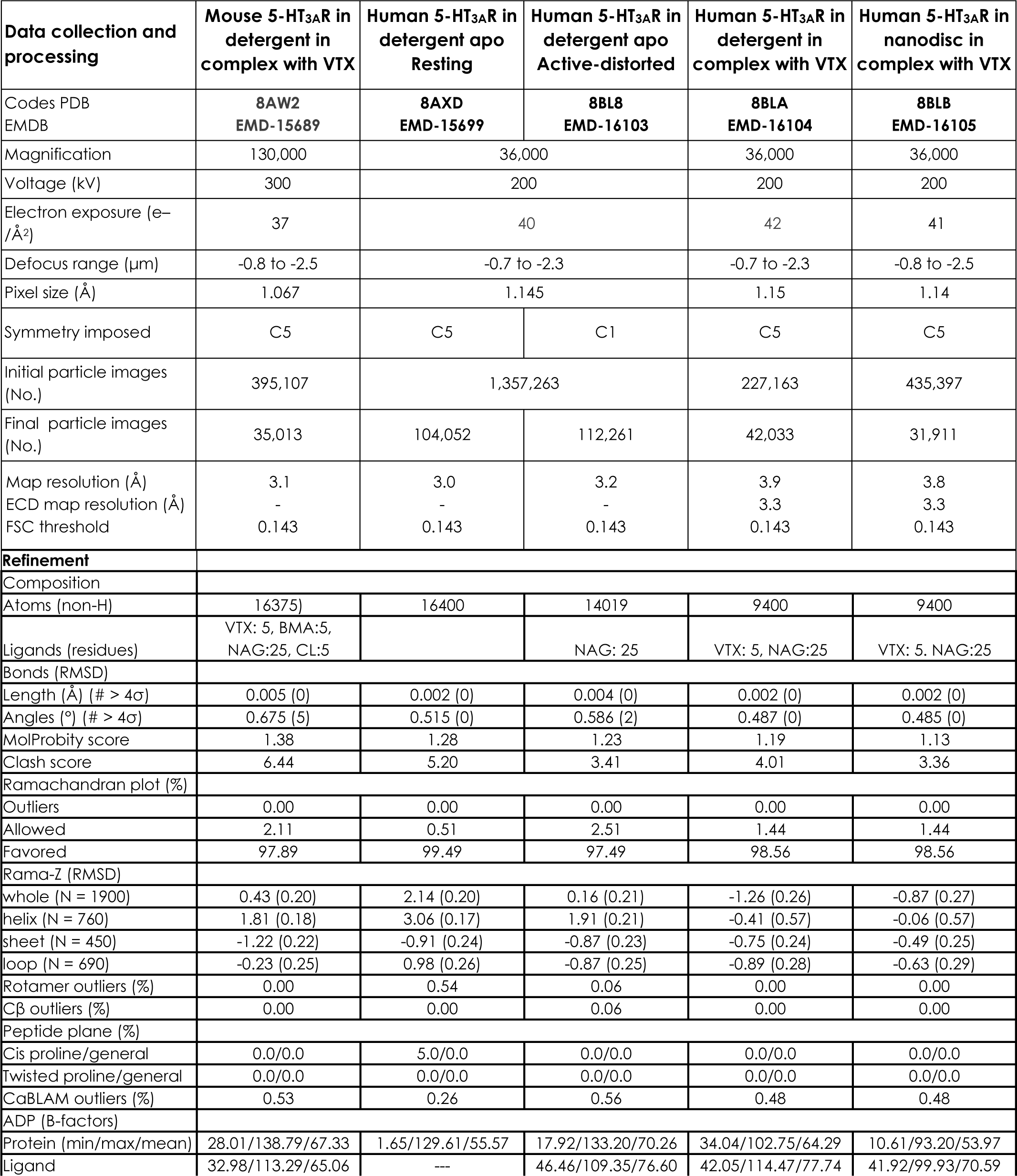
Cryo-EM data collection, refinement and validation statistics.

**Table SI4.** Structure-activity relationship of VTX analogs. The table shows the substitution patterns of the VTX scaffold for VTX and analogs and anolog agonist activity at WT and V202I mutant h5-HT3A receptors determined using the membrane depolarization assay at a concentration of 1 µM (Materials and Methods). For WT, the analog activity is nomalized to the response to 1 µM VTX. For the V202I mutant, the activity is normalized to the response to 10 µM 5-HT. The structure of the VTX scaffold with subsituent positions (R1-R8) are shown above the table.

| Ring |  |  |  |  |  |  |  |  | Activity (%) |  |
| --- | --- | --- | --- | --- | --- | --- | --- | --- | --- | --- |
|  | A | A | A | B | B | B | B | C | h5-HT<sub>3A</sub> WT | h5-HT<sub>3A</sub> V202I |
| Position | R1 | R2 | R3 | R4 | R5 | R6 | R7 | R8 |  |  |
| **Compound** |  |  |  |  |  |  |  |  |  |  |
| VTX | CH<sub>3</sub> | H | CH<sub>3</sub> | H | H | H | H | H | 100 ± 5 | 1 ± 1 |
| 4a | H | H | H | H | CH<sub>3</sub> | H | H | H | 1 ± 0 | 1 ± 1 |
| 4b | OCH<sub>3</sub> | H | H | H | CH<sub>3</sub> | H | H | H | 5 ± 3 | 0 ± 0 |
| 4c | Cl | H | H | H | CH<sub>3</sub> | H | H | H | 0 ± 0 | 1 ± 1 |
| 4d | F | H | H | H | CH<sub>3</sub> | H | H | H | 1 ± 1 | 0 ± 0 |
| 4e | CH<sub>3</sub> | H | H | H | CH<sub>3</sub> | H | H | H | 1 ± 0 | 2 ± 1 |
| 4f | H | H | H | H | H | H | H | H | 71 ± 12 | 4 ± 3 |
| 4g | OCH<sub>3</sub> | H | H | H | H | H | H | H | 101 ± 5 | 43 ± 11 |
| 4h | Cl | H | H | H | H | H | H | H | 95 ± 12 | 12 ± 3 |
| 4i | F | H | H | H | H | H | H | H | 80 ± 10 | 2 ± 1 |
| 4j | CH<sub>3</sub> | H | H | H | H | H | H | H | 95 ± 12 | 48 ± 9 |
| 4k | CF<sub>3</sub> | H | H | H | H | H | H | H | 67 ± 8 | 4 ± 3 |
| 4l | C(CH<sub>3</sub>)<sub>3</sub> | H | H | H | H | H | H | H | 57 ± 12 | 2 ± 1 |
| 5a | H | OCH<sub>3</sub> | H | H | H | H | H | H | 88 ± 16 | 3 ± 1 |
| 5b | H | CH<sub>3</sub> | H | H | H | H | H | H | 77 ± 14 | 1 ± 1 |
| 5c | H | H | OCH<sub>3</sub> | H | H | H | H | H | 93 ± 15 | 0 ± 0 |
| 5d | H | H | CH<sub>3</sub> | H | H | H | H | H | 100 ± 16 | 0 ± 0 |
| 5e | OCH<sub>3</sub> | OCH<sub>3</sub> | H | H | H | H | H | H | 98 ± 5 | 49 ± 12 |
| 5f | Cl | Cl | H | H | H | H | H | H | 51 ± 16 | 1 ± 1 |
| 5g | CH<sub>3</sub> | CH<sub>3</sub> | H | H | H | H | H | H | 94 ± 16 | 20 ± 4 |
| 5h | H | OCH<sub>3</sub> | OCH<sub>3</sub> | H | H | H | H | H | 91 ± 6 | 2 ± 1 |
| 5i | H | Cl | Cl | H | H | H | H | H | 71 ± 10 | 5 ± 3 |
| 5j | H | CH<sub>3</sub> | CH<sub>3</sub> | H | H | H | H | H | 85 ± 8 | 9 ± 6 |
| 5k | OCH<sub>3</sub> | H | OCH<sub>3</sub> | H | H | H | H | H | 87 ± 12 | 0 ± 0 |
| 5l | Cl | H | Cl | H | H | H | H | H | 87 ± 6 | 30 ± 3 |
| 5n | OCH<sub>3</sub> | H | Cl | H | H | H | H | H | 111 ± 14 | 61 ± 9 |
| 5o | CH<sub>3</sub> | H | OCH<sub>3</sub> | H | H | H | H | H | 104 ± 19 | 3 ± 1 |
| 5p | CH<sub>3</sub> | H | Cl | H | H | H | H | H | 96 ± 12 | 49 ± 5 |
| 6a | CH<sub>3</sub> | H | CH<sub>3</sub> | CH<sub>3</sub> | H | H | H | H | 33 ± 16 | 1 ± 0 |
| 6b | CH<sub>3</sub> | H | CH<sub>3</sub> | H | CH<sub>3</sub> | H | H | H | 0 ± 0 | 1 ± 0 |
| 6c | CH<sub>3</sub> | H | CH<sub>3</sub> | H | H | CH<sub>3</sub> | H | H | 16 ± 3 | 0 ± 0 |
| 6e | CH<sub>3</sub> | H | CH<sub>3</sub> | H | H | H | H | CH<sub>3</sub> | 52 ± 18 | 1 ± 1 |

